# ANTIPODE Provides a Global View of Cell Type Homology and Transcriptomic Divergence in the Developing Mammalian Brain

**DOI:** 10.1101/2025.10.18.683238

**Authors:** Matthew T. Schmitz, Jingwen W. Ding, Sara Nolbrant, Reed McMullen, Chang N. Kim, Bryan J. Pavlovic, Tomasz J. Nowakowski, Trygve E. Bakken, Chun Jimmie Ye, Alex A. Pollen

## Abstract

Diverse neurons and glia are generated in conserved spatial and temporal sequences during mammalian brain development. Divergence in gene regulatory networks can alter brain composition, scaling, timing, and function. However, resolving the identity, extent, and principles of gene regulatory divergence requires cellular-resolution surveys spanning brain regions and species and improved methods for defining cell type homologies. Here, we present ANTIPODE, a deep-learning variational inference framework that simultaneously integrates single-cell datasets, identifies homologous cell types, and parcellates differential expression across cell types, modules, and covariance. Applying ANTIPODE to a census of the whole developing macaque brain and a meta-atlas of human, macaque, and mouse brain development, we find broad conservation of initial neuron classes but widespread regulatory divergence within homologous types, shaped by genomic context, cell lineage, and developmental timing. Together, ANTIPODE provides a formalized and interpretable framework for cross-species single-cell analysis and reveals principles of gene regulatory divergence in mammalian brain evolution.

## Introduction

The structure of the mammalian brain follows a conserved bauplan more than 150 million years old.(Glenn Northcutt and Kaas, 1995; Striedter, 2004) Across the rostrocaudal extent of the neural tube, progenitor expansion is followed by neurogenesis and gliogenesis, however, this process proceeds at vastly different rates across species and among different regions of the developing brain.(Charvet et al., 2011; Qian et al., 2000; Workman et al., 2013) The orchestration of progenitor division and progeny differentiation yields the construction of brains that vary widely in shape, size, and function. Understanding the diversity and evolution of cell types in the developing brain is further complicated by the cyclical nature of progenitor renewal, spatiotemporal state gradients, and the cascade of quasi-parallel differentiation trajectories of cellular maturation originating from myriad distinct germinal zones.(Cadwell et al., 2019; Javed et al., 2025; Nowakowski et al., 2017; Puelles et al., 2013)

Interspecies comparisons of gene expression across homologous cell types could provide insights into the conserved and divergent properties of mammalian brain development. Recent analytical strategies now enable predicting homologous cell types from single-cell gene expression.(Haghverdi et al., 2018; Hie et al., 2024; Johansen and Quon, 2019; Korsunsky et al., 2019; Tarashansky et al., 2021; Welch et al., 2019; Xu et al., 2021) However, homology determination often relies on heuristics such as shared nearest neighbors and nonlinear data transformations and is particularly challenging across continuous developmental trajectories. (Colonna et al., 2024) Similarly, determination of cell type-specific gene expression divergence relies on homology and is sensitive to altered distributions in gene expression.(Zhou et al., 2019) Moreover, gene expression divergence can be partitioned into distinct modes, including alterations in cell type-specific gene expression, gene covariance, and gene module expression. These modes, however, are obscured by analytical strategies for differential expression that only consider a few cell types or brain regions in isolation.(Bakken et al., 2021; Harris et al., 2021; Ovens et al., 2021) We reasoned that a model explicitly accounting for categories of gene expression divergence and developmental trajectories applied to a multi-species whole brain dataset could simultaneously reveal cell type homology and putative mechanisms driving cell type evolution.

We introduce ANTIPODE, Ancestral Node Taxonomy Inference by Parcellation Of Differential Expression, a model that considers inferring cell type homology via integration and modeling cross-species differential expression as two sides of the same coin. Applying ANTIPODE to a census of rhesus macaque whole brain development and a meta-analysis of human and mouse brain development, we uncover profound conservation of the developmental Euarchontoglire transcriptomic bauplan and elucidate the spatiotemporal progression of the progenitors which build the structures of these divergent brains. In this process, we reveal the gene expression architecture underlying cross-species and cross-regional heterochrony and find transcriptomic correlates for the classic "later is larger" principle of vertebrate brain development.(Finlay and Darlington, 1995)

## Results

### Construction of a cross-species developmental brain meta-atlas

To systematically explore the evolution of developmental cell types across species, we constructed a meta-atlas by integrating data that we generated from the developing rhesus macaque whole brain (Figure 1a) and 13 datasets (see Methods, Supplementary table 1) totaling 1,854,767 cells from 420 10X genomics scRNA-seq ports. These samples span periods of neurogenesis across regions and species: post-conception day (PCD) 10-21 in mouse, PCD 40-110 in macaque and PCD 21-175 in human (Figure 1b). We uniformly reprocessed these datasets by obtaining raw reads and quantifying genes using Kallisto to mitigate alignment and reference-based artefacts. After processing steps, including ambient RNA, doublet, and non-brain cell type removal (Methods), we developed a unified cross-species cellular atlas.

**Figure 1.**
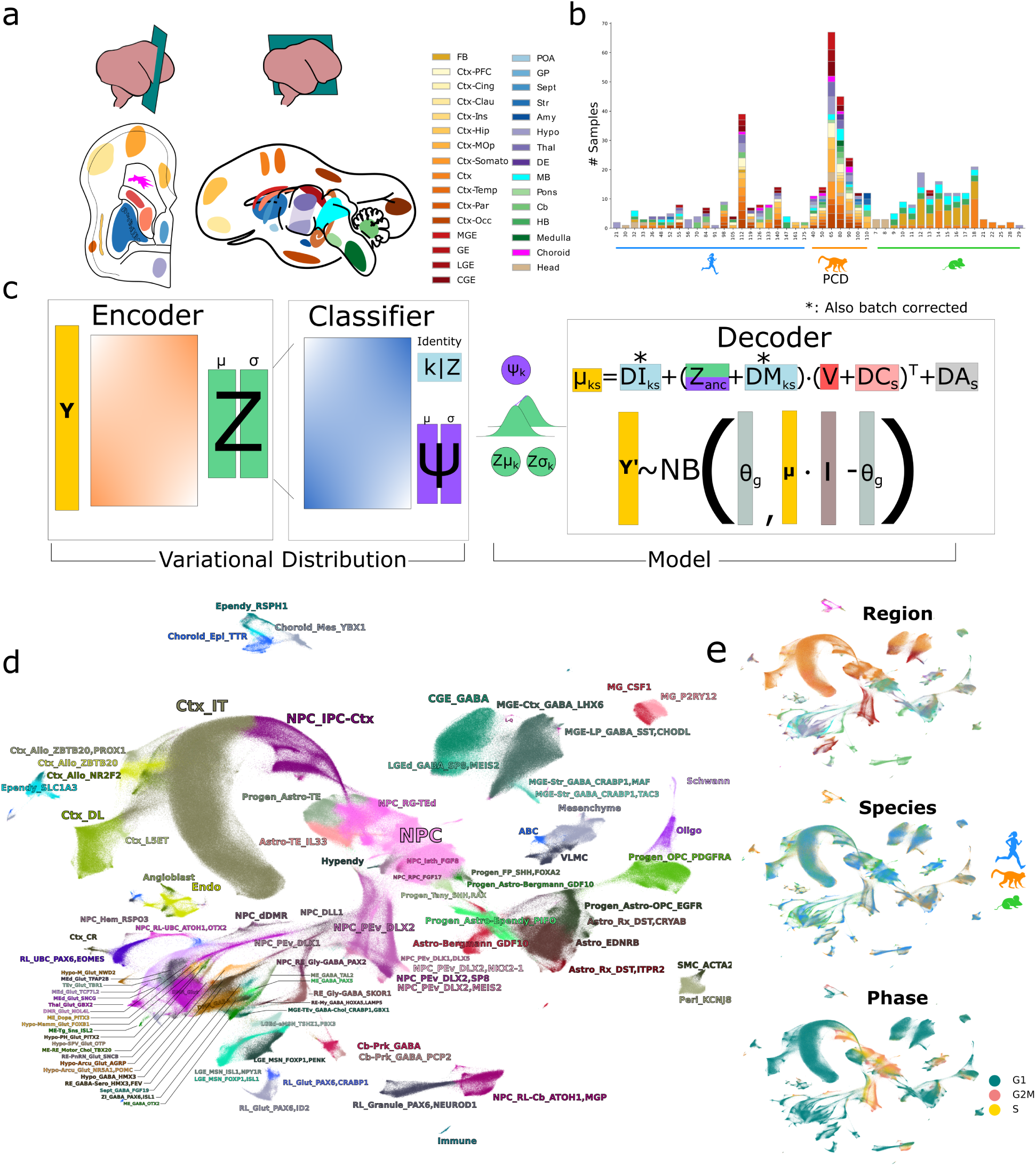
Construction of a cross-species developmental brain meta-atlas. **a,** Sampling strategy. Colored areas indicate the dissected structures in a PCD80 macaque brain; this regional color palette is used throughout. **b,** Composition of the atlas. Stacked bars show the number of samples ordered by post-conception day (PCD) per species, colored by region, see color legend in (a). **c,** Schematic of ANTIPODE. The encoder compresses gene counts to a latent variable Z; a classifier branch generates probability of each cluster k and ψ values; decoding is according to the equation shown with observed counts likelihood computed with the negative binomial distribution. **d,** Two-dimensional UMAP projection of the ANTIPODE latent space for all cells, colored by the 98 initial classes. **e,** UMAP projection colored by region (top, see color legend in (a)), species (middle) and by inferred cell-cycle phase (calculated by scanpy score_genes_cell_cycle) (bottom).

To identify homologous cell types, we introduce ANTIPODE, a novel variational inference model built using the pyro probabilistic programming language and inspired by scANVI, but designed specifically for simultaneous cross-species integration, de novo clustering and differential gene expression analysis (see full specification in Supplement 1).(Bingham et al., 2019; Xu et al., 2021) We modeled the expression of each gene in a cell type for a given species as a function of cell type-specific divergence, differential covariation, and differential module expression from a shared ancestral manifold of development (Figure 1c). To solve for these parameters while simultaneously clustering, we employed a maximum a posteriori variational autoencoder. This model applies the structural constraint that the discrete clusters and latent space generated by the encoder/classifier are decoded bilinearly using only the latent space, a single layer of weights and the differential expression parcels which are regularized by Laplace prior distributions (Figure 1b). This is in contrast to the typical nonlinear multi-layer perceptron decoder employed by most variational autoencoder models.(Lopez et al., 2018; Xu et al., 2021) In addition, the bilinear decoding of this model still allows removal of the linear effects of differentiation trajectories and secondary covariates like cell cycle phase, “regressing out” these effects in order to reconstruct observed expression values, which are modeled by a negative binomial distribution (Figure 1c).

We applied ANTIPODE to the developing brain meta-atlas to build a consensus taxonomy. Starting from 600 initialized clusters, the model identified 380 discrete state clusters. We next grouped these clusters into 98 initial classes, comprising the distinct progenitors and immediately post-mitotic transcriptional states in the developing brain (Figure 1e, Figure 2). (Schmitz et al., 2022) Clustering based on this method improved mixing between species and revealed normal distributions differential expression and covariance (Figure S2).

**Figure 2.**
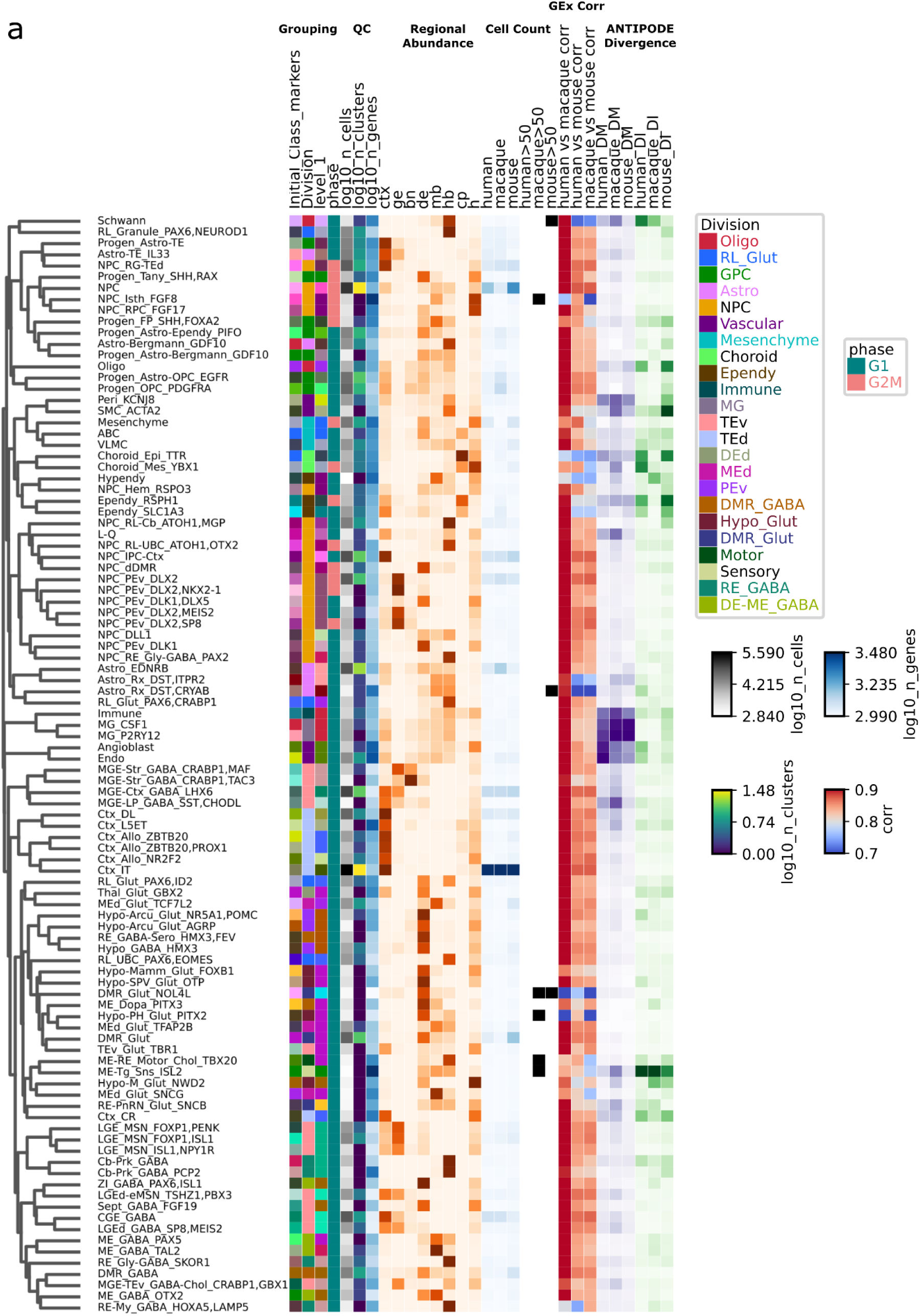
Taxonomy and ANTIPODE model architecture. Heat-map summary of the 98 initial classes. For each class (rows) we display (left-to-right): regional abundance across broad areas, total cell count, Spearman correlation of gene expression across pairs of species, mean absolute inter-species log fold change (“ANTIPODE divergence”). Row dendrogram shows Kendall correlation hierarchical clustering based on ANTIPODE latent space.

We next interpreted the cell identities according to enriched genes and anatomy (see Supplementary Table 2 for clusters, initial classes, and explanations). Briefly, non-neuronal cells were grouped into divisions: neuroepithelial/neural progenitor cells (NPCs); various types of HES6+ intermediate progenitor cells/neurons (IPC); glial progenitor cells (GPCs); vascular cells including angioblasts, endothelial cells, pericytes, smooth muscle cells (SMC); mesenchyme including arachnoid barrier cells (ABC), vascular/leptomeningeal cells, (VLMC); choroid plexus cells; oligodendrocytes; astrocytes including radial glia-like telencephalic astrocytes and non-telencephalic astrocytes; ependymal cells; microglia (MG); and other immune cells.(Rowitch, 2004) Newborn neuronal classes (TUBB3+) were similarly organized into divisions: ventral and dorsal telencephalic neurons (TEv, TEd); dorsal thalamic and mesencephalic neurons (DEd, MEd); secondary prosencephalic neurons (PEv) including the DLX+ hypothalamic fields; hypothalamic glutamatergic neurons (Hypo_Glut); diencephalic/mesencephalic GABAergic-like neurons (DE-ME_GABA); rhombencephalic GABAergic and/or glycinergic neurons; brainstem motor and sensory neuron classes; and heterogeneous groups termed diencephalic-mesencephalic-rhombencephalic (DMR) neurons (Figure 1c,2).(Bulfone et al., 1993; Moreno and González, 2011; Nieuwenhuys and Puelles, 2015) More mature neurons of arising from these classes were not captured, likely due depletion of larger neurons during whole cell dissociation. We provide supervised markers in the initial class name and conserved markers based on the minimum natural log fold change (LFC) in an initial class compared to the 99th percentile of expression in all other initial classes.(Figure 2,S3). Together, these initial classes, though likely not exhaustive, provide a foundation of homologous cell types for comparative analysis of Euarchontogliran development.

To link developing initial classes to putative terminal class derivatives in the adult brain, we used the list of identity-defining transcription factors (TFs) proposed in Yao et al. and performed pairwise Pearson correlation of cell type means to find them most correlated developmental and adult correspondences (Figure S4).(Yao et al., 2023) The supervised adult list qualitatively outperformed unsupervised lists of TFs filtered by variance and bimodality (data not shown), and we note that TF expression appeared more binary in adult cell types compared to developmental initial classes (Figure S5). In absence of whole-brain fate mapping data, we used developmental vs adult transcription factor correlations to assess the coverage of initial classes corresponding to adult subclasses. Fewer than 20 of 337 adult subclasses (mostly from medulla) have a developing initial class correlation less than 0.33 (greater than the 0.95 quantile of non-max correlations), supporting candidate linkages showing coverage of the precursors of most adult types in our atlas (Figure S4). Across the entire brain, among the initial classes we define, we do not find evidence of the absence of any initial class in any species, consistent with recent studies of telencephalic inhibitory neurons.(Corrigan et al., 2024) We do note, however, that these initial classes represent only a fraction of the adult diversity of mature cell types. For example, caudal ganglionic eminence-derived (CGE) cortical GABAergic interneuron initial classes appear uniform in prenatal development but diverge substantially into multiple subclasses (e.g., SNCG, PAX6, LAMP5, LAMP5/LHX6, VIP) and many more types in the adult brain. Similarly, undifferentiated developmental classes likely mask further developmental diversity. This is especially true for classes like DMR_Glut or DMR_GABA, which contain cells from multiple regions of the ventral brainstem, and appear to correspond roughly to the diverse neuron types previously described as forming “splatter” clusters in the adult human brain.(Siletti et al., 2023)

### ANTIPODE as a cross-species integration method and differential expression model

To provide additional validation of the ANTIPODE model as an integration method, we tested its capacity for batch correction and preservation of biological variation on two additional cross-species test datasets. One dataset included snRNAseq data from many human cortical areas along with many primates from a single area, and the other included adult retina snRNAseq from 13 highly diverged mammals.(Hahn et al., 2023; Jorstad et al., 2023b, 2023a) ANTIPODE demonstrated superior performance in cross-species integration tasks compared to established methods such as Harmony and SCANORAMA, and performed comparably to SCVI, despite these other methods utilizing much stronger nonlinear integration approaches (Figure 3b-h, S6). (Hie et al., 2024; Korsunsky et al., 2019; Lopez et al., 2018) Crucially, ANTIPODE also simultaneously provides clustering and differential gene expression analysis during its fitting procedure, offering practical advantages over other models requiring sequential analytical steps. As such, ANTIPODE was effective at the reconstruction of log expression values, with an R² value of 0.90 for log expression means across all species, types and genes (Figure S2). On the other hand, because ANTIPODE models real differential expression in log fold change space, it is likely not suitable for nonlinear integration tasks such as integration of single-cell RNA data with single nucleus or spatial transcriptomics data.

**Figure 3.**
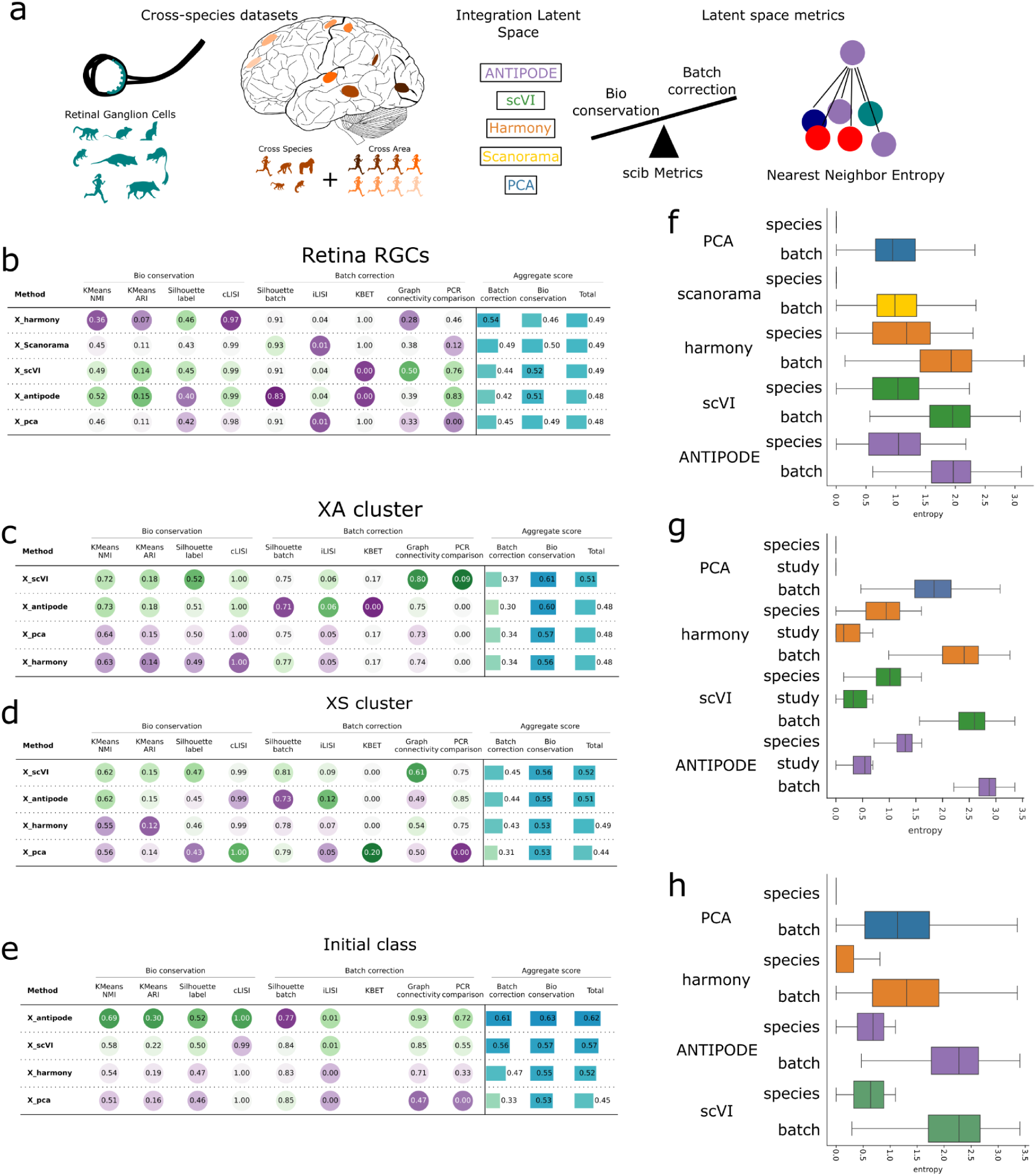
Benchmarking ANTIPODE against existing cross-species integration methods. **a,** Schematic detailing the benchmarking strategy for evaluating ANTIPODE as an integration method. **b–e,** Bubble plots summarizing integration metrics calculated by scib for: (*b*) mammalian retinal ganglion cells from Hahn et al.,, (*c,d*) cortical cross-areal (XA) and cross-species (XS) clusters from combined Jorstad et al. and Jorstad et al.. (*f*) whole developing brain initial classes from the current study **f–h,** Aggregated integration space k-nearest neighbor classification entropy for each method’s latent space calculated for (*f*) mammalian retinal ganglion cells from Hahn et al.,(*g*) combined cortical cross-areal (XA) and cross-species (XS) clusters from combined Jorstad et al. and Jorstad et al., (*h*) whole developing brain initial classes from the current study. Boxplots show median and inter-quartile range.

### Principles of evolutionary divergence in developmental gene expression

Approximately 71% of genes exhibited at least a two-fold expression change between at least one pair of species, violating typical assumptions underlying null models for differential expression (DE) analysis (Figure 4a). Moreover, continuous cellular identity changes during development make instantaneous gene expression dispersion estimates challenging. Therefore, instead of focusing on specific differentially expressed genes, we examined broad evolutionary trends across developmental gene expression patterns by jointly modeling gene expression across developmental states and species-specific divergence by differential expression categories (Figure 4b). We used ANTIPODE’s learned parameters to separate 4 categories of differential expression (DE), where differential-by-all (DA) is a species intercept representing DE across all cell types, differential-by-identity (DI) representing DE in a single-cell type, differential-by-module (DM) representing DE of an entire module of genes in a single type and differential-by-coexpression (DC) representing differential module membership of genes across species (Figure 4b). We found that in our model, the effects of DM and DC are correlated, as these are multiplied together, with Spearman correlations of effects within taxons around 0.7, while the correlation of either to DI is much lower (Figure S7d).

**Figure 4.**
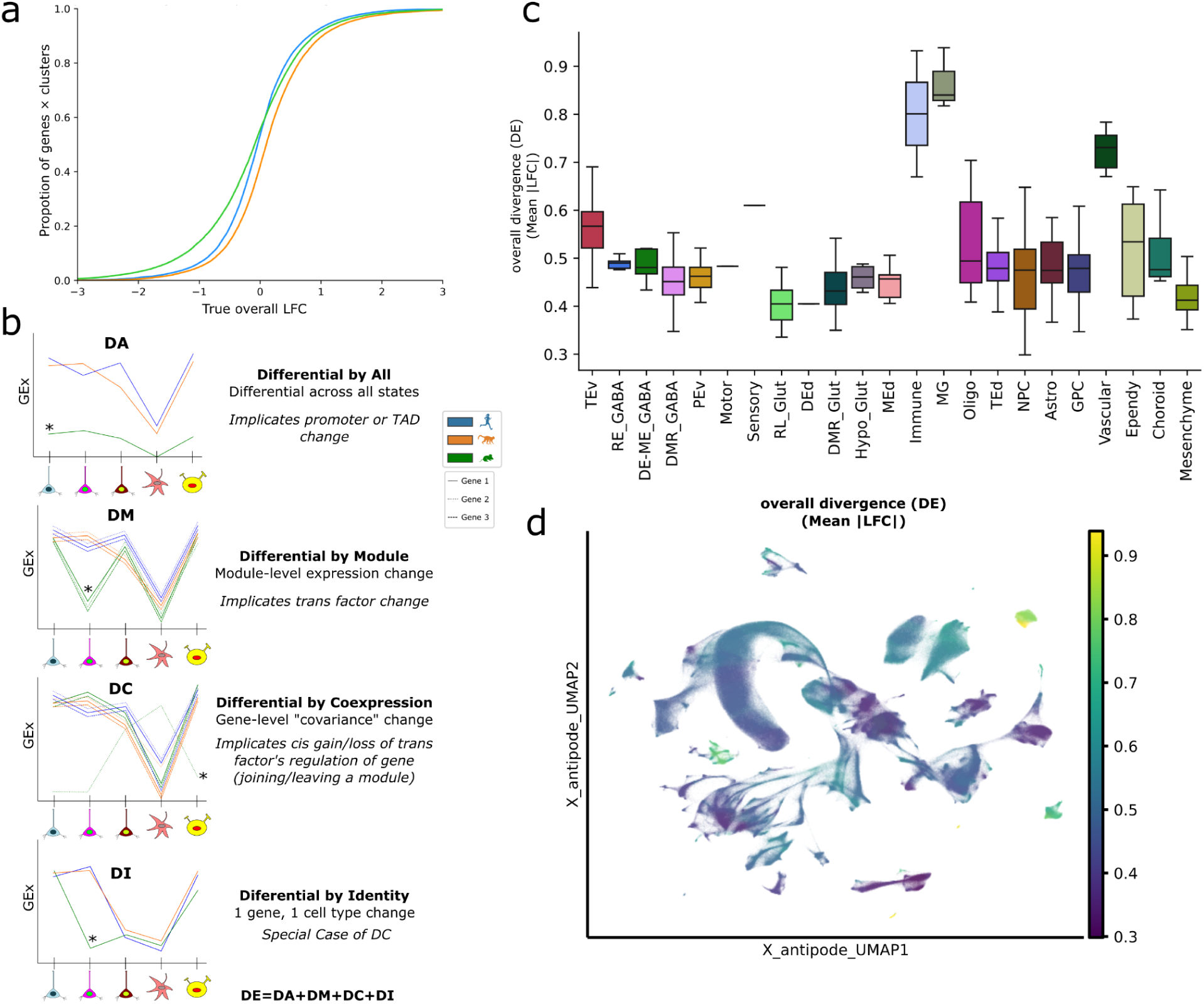
Modes and landscape of evolutionary differential expression. **a,** Cumulative distribution showing proportion of genes by their per species log fold change. 71.6% exceed |log2FC > 1 in at least one pairwise species comparison (based on raw log expression, not ANTIPODE fit). **b,** Conceptual schematic of four categories of differential expression learned by ANTIPODE: DA (differential-by-all states intercept), DM (by-module), DC (by-co-expression) and DI (by-identity intercept). **c,** Boxplot showing median and quartile divergence (overall DE) for each ANTIPODE cluster, grouped by division. **d,** ANTIPODE UMAP colored by each cluster’s summary divergence (mean |LFC for all categories combined).

We observed that microglia showed the highest divergence among brain cell types across species, with ependymal cells also having higher overall DE divergence, aligning with prior reports, while other endogenous glia like astrocytes and oligodendrocytes diverge similarly to most neurons despite having higher divergence in adult cortex (Figure 4c,d).(Jorstad et al., 2023a) Intriguingly, the small number of brain-exogenous immune cells captured also displayed similarly high divergence, suggesting rapid microglial evolution may be attributed more to their immune/myeloid lineage rather than to brain-specific selective pressures. We also found that as expected, gene expression divergence tends to increase in neurons relative to progenitors (Figure 4c,d).

We next looked for associations between genomic context and the strength of divergence across categories. The strongest association was between gene length and DA (slope LFC/log length), with large genes showing a strong reduction in DA compared to smaller genes (Figure 5a,b). Non-TF genes and TFs not expressed in both developing initial and mature classes had similar divergences across categories, and were the most DA. We also found that shared TFs for non-neuronal cells displayed greater divergence in general than neuronal TFs in DC and DM, while neurons tended to have higher DI, possibly due to their diversity and relatively fewer clusters per initial class (Figure 5c). Additionally, genes located in conserved syntenic genomic regions exhibited coordinated, modular differential expression, whereas genes in non-syntenic regions tended to be broadly and diffusely differentially expressed across many/all cell types (Figure 5d).

**Figure 5.**
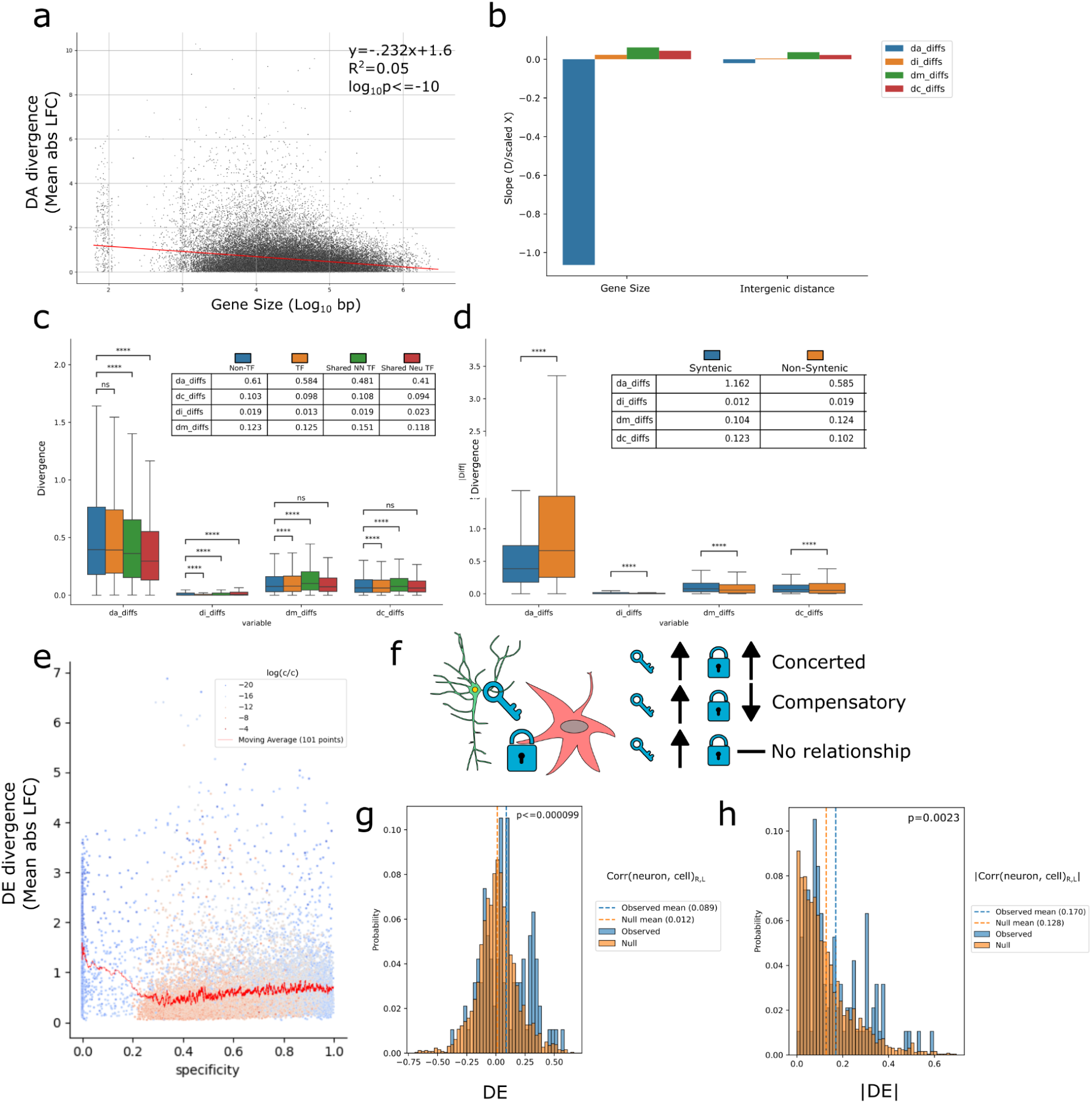
Gene expression evolution in context. **a,** Gene size vs DA for all genes. Line of best fit calculated by scipy linregress. **b,** Best fit line slopes between divergence categories vs 0-1 scaled log10 gene size and 0-1 scaled log10 intergenic distance. **c,** Divergence of DA, DI, DM and DC sets split by gene class (Shared Neu TF refers to TFs with shared expression in development and adult in neurons, Shared NN TF refers to TFs with shared expression in development and adult in nonneurons, TFs refers to TFs not shared between development and adult, Non-TF refers to all other genes not in one of the other categories). P values are Bonferroni corrected two sided Mann-Whitney test, from python’s statannotations package * p<0.05, ** p< 0.01, *** p<0.001, **** p<0.0001. **d,** Divergence of DA, DI, DM and DC sets split by conserved synteny. P value annotations are the same as **c. e,** Summary divergence versus specificity (tau, 0-1 score where 1 represents gene is expressed only by one type) colored by log gene mean expression value. **f,** Schematic depicting evolutionary implications of paired neuropeptide and receptor gene expression divergence in sender and recipient initial classes, respectively. Neuropeptides are represented by keys and receptors are represented by locks. **g–h,** Histogram showing observed ligand–receptor correlations (blue) are more concordant in direction (*g*) and magnitude (*h*) than 10 000 shuffled pairs (orange); vertical lines indicate mean values.

Given the simplified pairing of neuropeptides and their receptors (often one-to-one gene pairs), we hypothesized these pairings could serve as a test case for the extent of coordinated gene expression during evolution. Indeed, differential expression analysis revealed ligand and receptor genes frequently evolving cooperatively, showing directional changes that were significantly more often in the same direction (p < 0.000099), and coincidentally stronger (p = 0.0023) than expected by chance (Figure 5f-h, S8).

### Spatiotemporal dynamics of progenitor states and neurogenic timing

The timing of mammalian brain development varies globally across species and sequentially across regions. For example in the pallium of the mouse the vast majority of neurogenesis occurs between PCD 10.5 to 18.5 while birth is at day 21, in the macaque occurs between PCD 40 and 100 while birth is around day 166, and in the human occurs between PCD 42 and 161 while birth is around day 280.(Di Bella et al., 2021; Rakic, 2002; Vanderhaeghen and Polleux, 2023) Meanwhile the neurogenic window of different brain structures shows dramatic heterochrony, where in the mouse: the ventral midbrain spans PCD10.5-14.5; the dorsal midbrain spans PCD12.5-16.5; the ventral-derived pontine neurons and Purkinje cells spans PCD9.5-11.5; rhombic lip-derived pons, cochlear nuclei and cerebellar granule cells span PCD12.5 to postnatal stages, and strikingly, developmental neurogenesis of olfactory bulb granule neurons continues well after birth.(Gritti et al., 2002; Hirata et al., 2021) Thus, we next sought to examine the relationship between gene expression and global and stereotypical aspects of developmental timing.

We developed a Bayesian model to examine the temporal dynamics of progenitor states, modeling their abundances across developmental time with asymmetric split Gaussian distributions (Figure 6a). From these models, we calculated the "lateness", defined as the temporal center-of-mass (COM) of progenitor abundance. As expected, the temporal COM of astrocyte and OPC progenitor clusters followed that of neural progenitors across regions.

**Figure 6.**
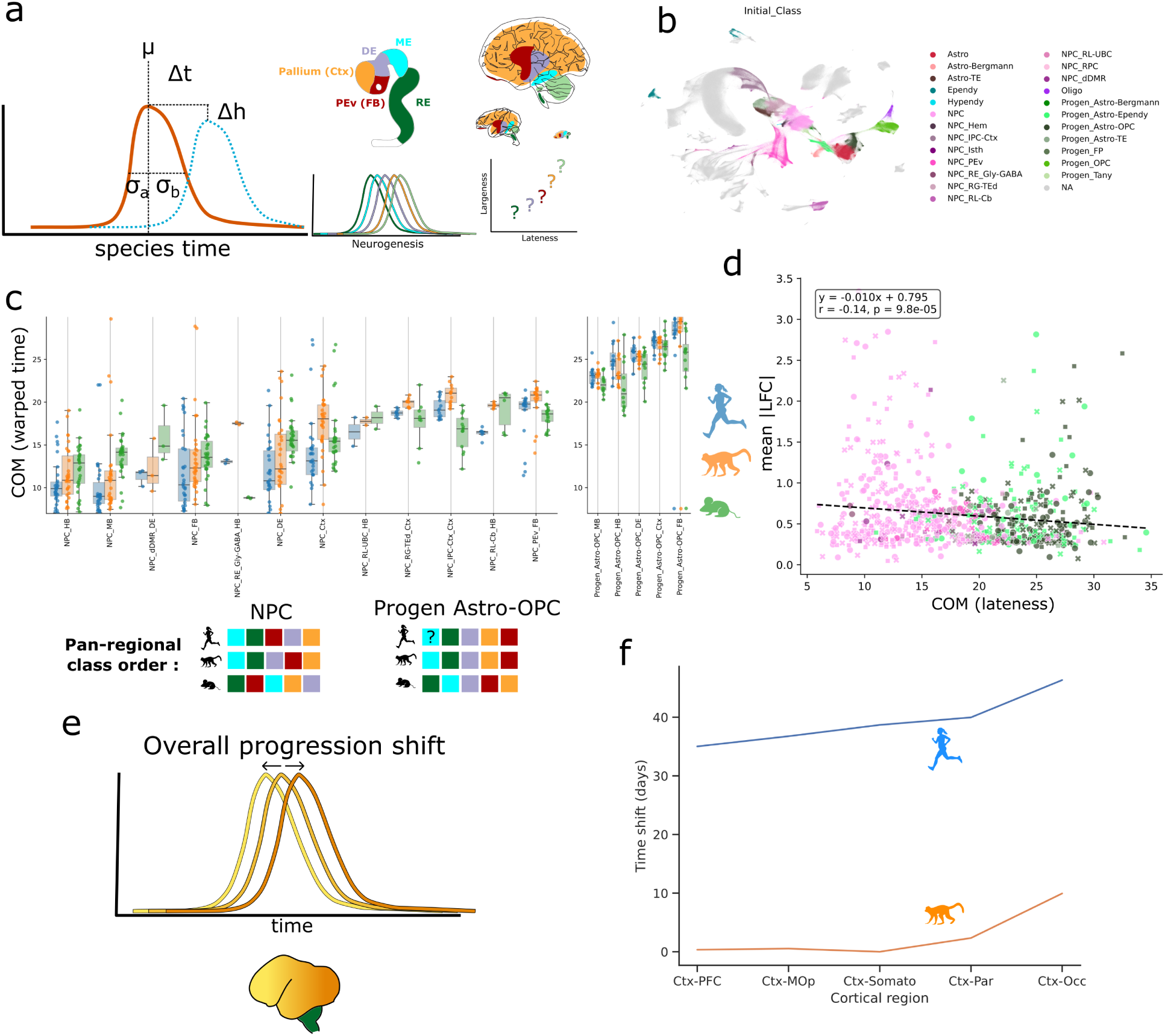
A Bayesian model of progenitor-state progression across species and regions. **a,** Progenitor timing model schematic. Progenitor abundances are fitted with asymmetric split-Gaussian curves with parameters: μ, σₐ, σᵦ, Δ*t*, Δ*h*; right inset is a cartoon depicting that larger structures tend to be later-developing. **b,** UMAP of progenitor classes used for timing model, colored by initial class. Abbreviations used throughout: (HB/RE: hindbrain/rhombencephalon, MB/ME: Midbrain/Mesencephalon, DE: Diencephalon, FB: Forebrain i.e. non-pallial, non-diencephalic prosencephalon, Ctx: Cortex i.e. pallium) **c,** Estimated centre-of-mass (COM) lateness for distinct progenitor state across regions (points colored by species) with NPC classes on the left and Astro-OPC progenitors on the right (these are the progenitor groupings that are analogous across all regions). Boxplots show median and quartile values. Boxplot including all values can be found in Supplementary Figure 9. **d,** Scatter of overall mean |logFC DE versus stereotypical COM for each progenitor state; color encodes the initial class. **e,** Schematic summarizing the known acceleration rostral-caudal spatiotemporal neurogenic gradient. **f,** Relative acceleration of cortical progenitor progression fit by progenitor state model using only cortical regions along the rostral-caudal axis for human and macaque.

Among neural progenitors, we found regional differences consistent with the "later-is-larger" developmental principle: cortex, ganglionic eminence, and cerebellum had late COMs, whereas neuroepithelial states, ventral midbrain and myelencephalon showed early COMs (Figure 6c). (Finlay and Darlington, 1995) Using the ordering of progenitor states across time, we hypothesized that gene expression would become increasingly divergent later in development. We however found the contrary, a weak negative relationship between a state’s lateness and overall expression divergence (DE) (Figure 6d). Finally, in a second model which included an overall regional shift term, fitting with only cortical progenitors from prefrontal, motor, somatosensory, parietal and occipital cortex, progenitor state progression was accelerated in macaque relative to human and rostral cortex relative to caudal cortex (Figure 6e-f). This aligns with known neurogenic gradients and previous observations of accelerated primate visual cortex progenitor progression relative to the frontal lobe.(Nowakowski et al., 2017)

### Genes associated with developmental timing in progenitors

We then analyzed gene expression relationships with progenitor timing across species and regions. We considered three categories of timing differences: first "global lateness" captures the absolute timing of development across regions and species (e.g. the ordering mouse medulla, mouse cortex, macaque medulla, human medulla, macaque cortex, human cortex), second "stereotypical lateness" captures the conserved progression of states (e.g. the ordering medulla, diencephalon, cortex which is stereotypical to all species), and third, "differential lateness" which estimates shifts in the lateness of a state relative to that state’s stereotypical lateness. We scored genes against these metrics by calculating the correlation of a gene’s expression in states with those states’ lateness metrics (Figure 7a, S9f). Stereotypical lateness had the most genes with strong signal, followed by global lateness, while differential lateness’ signal was much more limited (Figure 7b). We also found that the magnitude of stereotypical lateness/earliness was most related to overall DI amongst progenitor types (Figure 7c).

**Figure 7.**
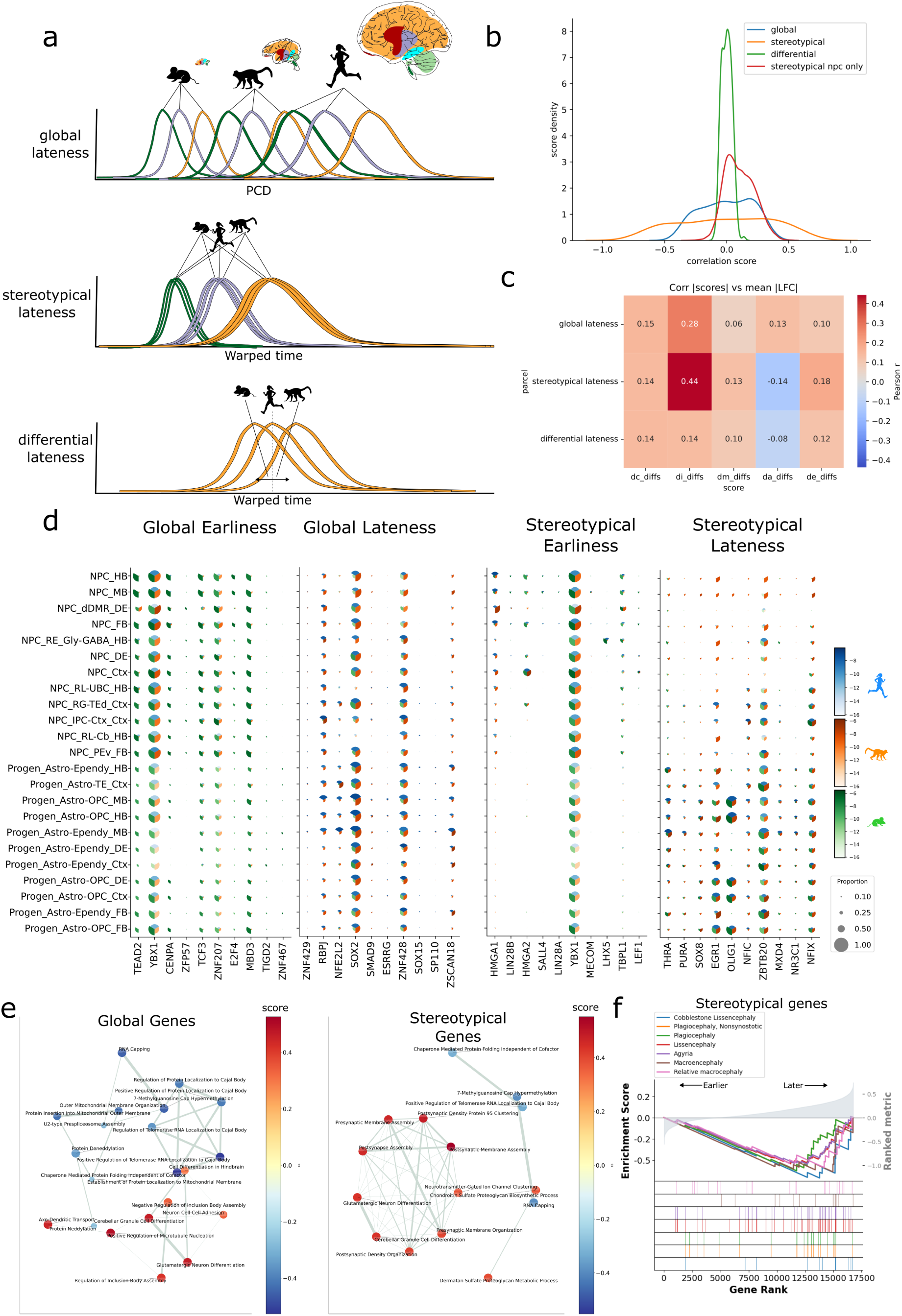
Gene programs linked to early versus late neurogenesis. **a**, Illustration of three lateness metrics: global, stereotypical and differential. **b**, Histogram of gene expression correlation scores with each progenitor lateness fit values **c,** Heatmap showing the correlation between the magnitude of each score category with the magnitude of each differential expression parcel’s overall divergence. **d**, Dot-plot of top genes correlated with each lateness metric (dot size = fraction expressing; color = log counts/count). **e**, Enrichment networks for genes associated with stereotypical lateness (left) and global lateness (right). Terms represent the top 15 positive and negative terms with resampling pvalue less than FDR<0.05. Edge width represents proportion of genes overlapping in gene sets, node color represents the mean lateness score for the genes in associated with that term. **f**, Gene set enrichment analysis (using gseapy prerank) of stereotypical lateness genes shows enrichment for neuromorphological disorder gene sets; curves plot directional enrichment score against ranked gene list.

Genes related to global lateness often displayed constitutive differences between species, many explained by pan-state DA in ANTIPODE, as expected, with the highest scored genes also showing temporal progression. Among these top global lateness genes, upregulation of transcription factors TEAD2, YBX1, TP53 was associated with mouse and global earliness while SOX2 was associated with global lateness and thus human (Figure 7d). There were no significant brain size malformation-related associations with global or differential lateness, but gene ontology clearly shows dynamics in mitochondrial respiration, splicing, translation and differentiation identity genes (Figure 7e-f).

We next examined the properties of genes related to stereotypical timing across species. Genes associated with smaller, early-developing brain regions (stereotypical earliness) included known pluripotency, proliferative factors, and oncogenes (e.g., LIN28A/B, IGF2BP1/2, HMGA1/2, SNRPE/F/G/A1, SALL4), whereas late-developing regions (stereotypical lateness-associated) expressed genes previously linked to growth restriction (e.g., NFIA/B/C/X, CEBPD, NDRG2, EGR1, PURA) (Figure 7d).(Nowakowski et al., 2017; Pollen et al., 2014) Many of these genes have been linked to timing of cortical development in previous studies, and our findings extend this timing relationship to developmental sequences across regions that are shared across species.(Nishino et al., 2013; Pollen et al., 2019) To add to this point, all 15 genes associated with macroencephaly by DisGeNET including AKT1/3, PTEN, PIK3CA/B/D/G and MTOR were positively correlated with stereotypical lateness. Genes associated with microcephaly, lissencephaly and agyria were all significantly associated with lateness as well (Figure 7f). We found a similar set of genes in the cortex area-only model, with notable additions of FZD3 as a top 5 gene related to cortical earliness, while NR2F1 and noted outer radial glia marker FGFR3 are the top genes related to cortical lateness (Supplementary Table 3) (Schaberg et al., 2022).

Finally, we examined the properties of genes with shifts in timing between species across regions, relative to stereotypical timing. Perhaps consistent with continuous, mosaic evolutionary modifications of brain structures, there was minimal coherence in the functional annotations of genes associated with "differential lateness", and most of these genes are associated with regional identity (e.g. HOX genes). While we cannot exclude that an increased temporal resolution of sampling would resolve convergent properties of altered regional timing, the limited coherence in this analysis suggests that diverse genetic mechanisms drive shifts across regions and species.

## Discussion

Here we built a model of transcriptomic shifts across evolution, ANTIPODE, to create a unified taxonomy of the initial classes across the developing Euarchontoglire brain. The model performed at least as well as other widely-used integration methods showing that cross-species gene expression divergence, despite presumed complexity, can be captured by an interpretable model. ANTIPODE achieved this integration as a bilinear model with interpretable parameters that correspond to intuitive categories of gene expression divergence, while performing simultaneous de novo clustering. From the three species developing brain meta atlas we constructed, ANTIPODE detected clusters that we grouped into 98 initial classes, which appear to be the precursors of most of the subclasses seen in the adult brain. In our data, which is enriched for progenitors and newborn neurons and depleted for mature neurons due to whole cell dissociation, we did not observe comparable neuronal diversity in any developing region to the adult brain, strongly suggesting that the outstanding diversity of neuronal types arises post-mitotically across all brain regions as neurons settle in terminal niches and crystallize into stable adult types.(Cheng et al., 2022; Fishell and Kepecs, 2020; Kim et al., 2020)

Using ANTIPODE as a holistic differential expression method, despite the profound conservation of cell states across species, we uncovered several patterns of gene expression evolution across species. We found that the largest share of gene expression divergence could be attributed to DA, changes across all/many types, while the other categories of differential expression were complex and entangled. Additionally because of the prevalence of DA, DM and DC changes which represent gene expression shifts across multiple types, we stress the importance of holistic differential expression accounting across many cell types and regions, lest one assign shifts in expression across many cell types as a cell type-specific change.

The prevalence of differential expression, with more than two-thirds of genes having a fold change of 2 or greater for a given type between any two of our three species, also suggests that the hunt for consequential transcriptomic changes across species underlying particular physiological traits may be “searching for a needle in a needlestack” where many changes push a quantitative trait in opposing directions to reach a new equilibrium. We acknowledge that post-transcriptional regulation may buffer many of these changes such that they have small effects on the proteome. Amidst such a haystack of gene expression divergence, however, there must also be cases of oppositional changes in expression by genes in the same pathway to compensate, and still other changes which are incompletely opposed and contribute to the observed morphological and functional differences across species’ brains.(True and Haag, 2001) Indeed, in the limited case of neuropeptide and receptor evolution, even though we could not filter our sender and receiver types by peptide-connectomics, we saw a significant coincidence of both ligands and receptors shifting together and in the same direction.

Finally, with a vantage point of development in three species with highly different developmental rates, we also examined the evolution of progenitor state composition through neurogenesis. Using a simplified model of progenitor state composition over time, we estimated the stereotypical ordering of progenitor states across regions and species, and how the different species differ in the ordering and occupancy of states through development. As it has been shown that in general gene expression becomes more divergent across species as cells in the developing embryo mature (Cardoso-Moreira et al., 2019), it has also been assumed that stem cells with higher potency are more constrained due to the potential of pleiotropic effects propagating in all descendant lineages. Counter to this expectation, while expression divergence increased in many post-mitotic states, it was essentially constant (slightly decreasing) as progenitors progressed from higher (neuroepithelium and NPC) to lower potency states (glial progenitors). This suggests that the constraint on gene expression may not be strictly due to the pleiotropy of multiple descendant cell types, and future perturbation studies will be able to answer how gene expression changes in progenitors of various states and potency affect the overall structure and function of the brain.

We identified genes with expression that correlates with progenitor lateness. Because developmental lateness and structure size are closely linked, these genes likely also include regulators of mosaic regional expansion.(Finlay and Darlington, 1995; Herculano-Houzel, 2012) We document a surprising, but not unprecedented mix of genes associated with lateness, with many genes expected to drive both expansion and attenuation of growth in NPCs. Here we see the power of a global view of brain development, where we saw the upregulation of PI3K and mTOR pathways reported in the context of cortical expansion as key levers for progenitor proliferation across many regions.(Andrews et al., 2020; Nowakowski et al., 2017; Pollen et al., 2019)

While the present dataset captures the majority of canonical neurogenesis across the three species, it has relatively low temporal resolution in primates and lacks coverage of late-stage gliogenesis, which extends significantly beyond birth. Thus, estimates of late glial progenitor dynamics are likely incomplete, and could be greatly improved as larger single nuclei datasets spanning more species, timepoints and including molecular recording technologies like neuronal birthdating or lineage barcoding are created and collated.(Bandler et al., 2022) Study of diverse vertebrates with dramatic regional expansions like the primate cortex, the cyprinid vagal lobe, the mormyrid cerebellar valvula, or various other instances of mosaic regional expansion in vertebrate lineages will give better opportunity to elucidate the degree to which independent expansions converge on similar pathways or whether it is idiosyncratic.(Striedter and Northcutt, 2019) Soon, it will likely be possible to look with much higher resolution to determine subtle shifts in developmental timing and to conclusively determine genes expression signatures that might be convergently responsible for shifts toward earliness or lateness across species outside of those already associated with global/stereotypical lateness. ANTIPODE gives us a window into this point, as genes more strongly related to stereotypical earliness/lateness tended to be much more DI, lending support to the idea that species may control allometric shifts by mosaic changes in gene regulation.

ANTIPODE represents a framework for modeling gene-expression divergence across species. The current implementation returns maximum a posteriori parameter estimates rather than full posteriors, so we cannot yet provide credible intervals or formal significance for individual parameters. While the architecture admits phylogenetic structure, our attempts to include ancestral nodes led to clustering collapse, indicating the need for more stable variational families or priors. In addition, the structure of the model makes it difficult to directly control clustering resolution and rare cell types in the adult brain. Finally, the bilinear decoding that makes parameter interpretation straightforward limits applicability to strongly nonlinear integration tasks (e.g., single-cell with single-nucleus or spatial); constrained nonlinear corrections are a plausible extension.

ANTIPODE brings integration, taxonomy, and differential expression into a single framework, converting discrete steps that often undermine one another into a coherent whole in which the latent space is anchored in cell-state structure and integration is accomplished by modeling differential expression. Applied to human, macaque, and mouse development, it reveals broad conservation of initial classes alongside pervasive, context-dependent divergence shaped by genomic architecture, lineage, and developmental timing. We identified coordinated evolution in ligand–receptor pairs and link progenitor state to developmental rates. In this work we have sought to zoom out and view species- or region-specific transcriptomic landscapes as parts of a larger global developing brain, and we foresee that biologically inspired models like ANTIPODE can shift analytical focus to a more universal view of the developing vertebrate brain.

## Supplementary Figures

**Supplementary Figure S1.**
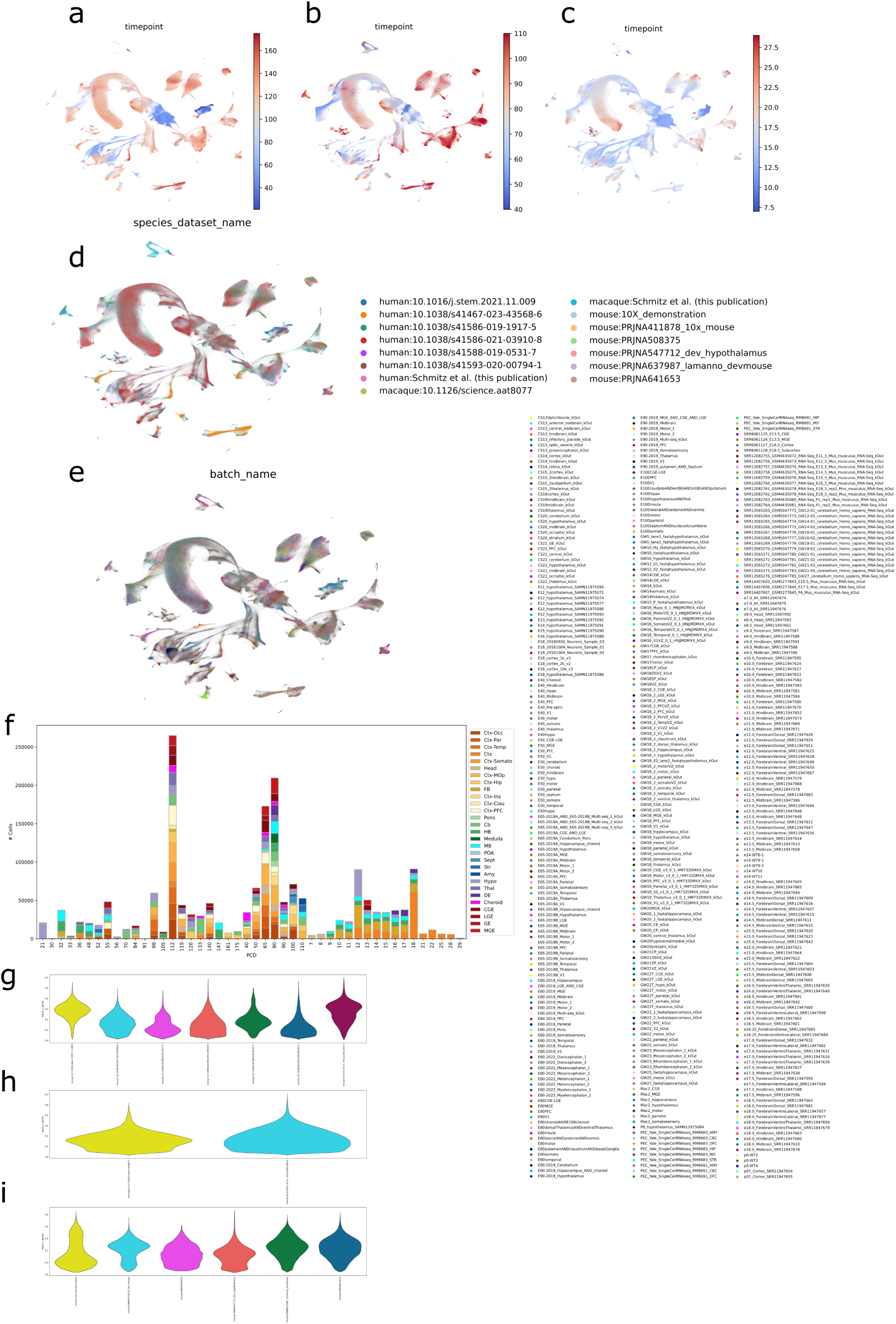
Dataset composition, batch structure and basic QC. **a–c,** ANTIPODE UMAP colored by post-conception day (PCD) for human (**a**), macaque (**b**) and mouse (**c**) reveal continuous developmental trajectories that align across species after integration. **d,** UMAP colored by the 15 10X scRNA-seq datasets showing that biological structure, not dataset of origin, dominates the embedding. **e,** The same UMAP colored by individual library-preparation batches. **f,** Stacked-bar plot of the 1,854,767 high-quality cells grouped by species, structure (colors) and PCD demonstrates broad temporal and anatomical coverage. **g–i,** Violin plots of log10 UMI counts per cell across datasets for human **(g)**, macaque **(h)** and mouse **(i)**.

**Supplementary Figure S2.**
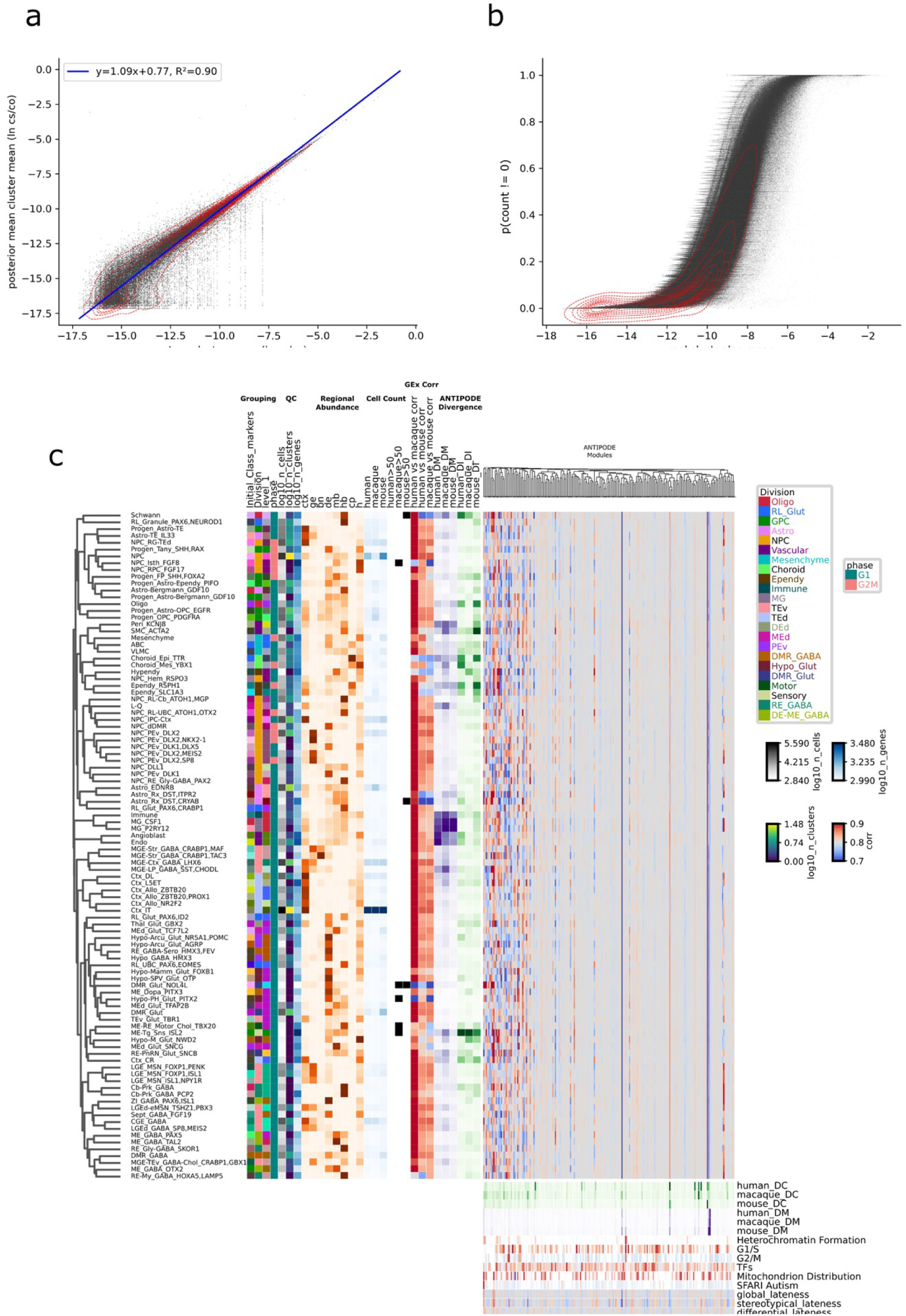
Goodness-of-fit for ANTIPODE gene-specific parameters. **a,** Reconstructed (posterior) mean log-expression (log counts/count) for every gene-cluster pair versus its empirical mean (black points; red density contours). **b,** Empirical zero probability (p(count = 0)) versus empirical log-mean expression suggest a small degree of zero-inflation (left-right shift of the curve). **c,** Full heat-map summary of the 98 initial classes from figure 2. For each class (rows) we display (left-to-right): regional abundance across broad areas, total cell count, Spearman correlation of gene expression across pairs of species, mean absolute inter-species log fold change (“ANTIPODE divergence”), membership in 200 ANTIPODE gene modules. The bottom section shows divergence by DC and DM. Gene set enrichments are colored by gseapy prerank normalized enrichment scores of each ANTIPODE component, with color alpha equal to 1 – significance q value. Row and column dendrogram shows Kendall correlation hierarchical clustering by ANTIPODE latent space.

**Supplementary Figure S3.**
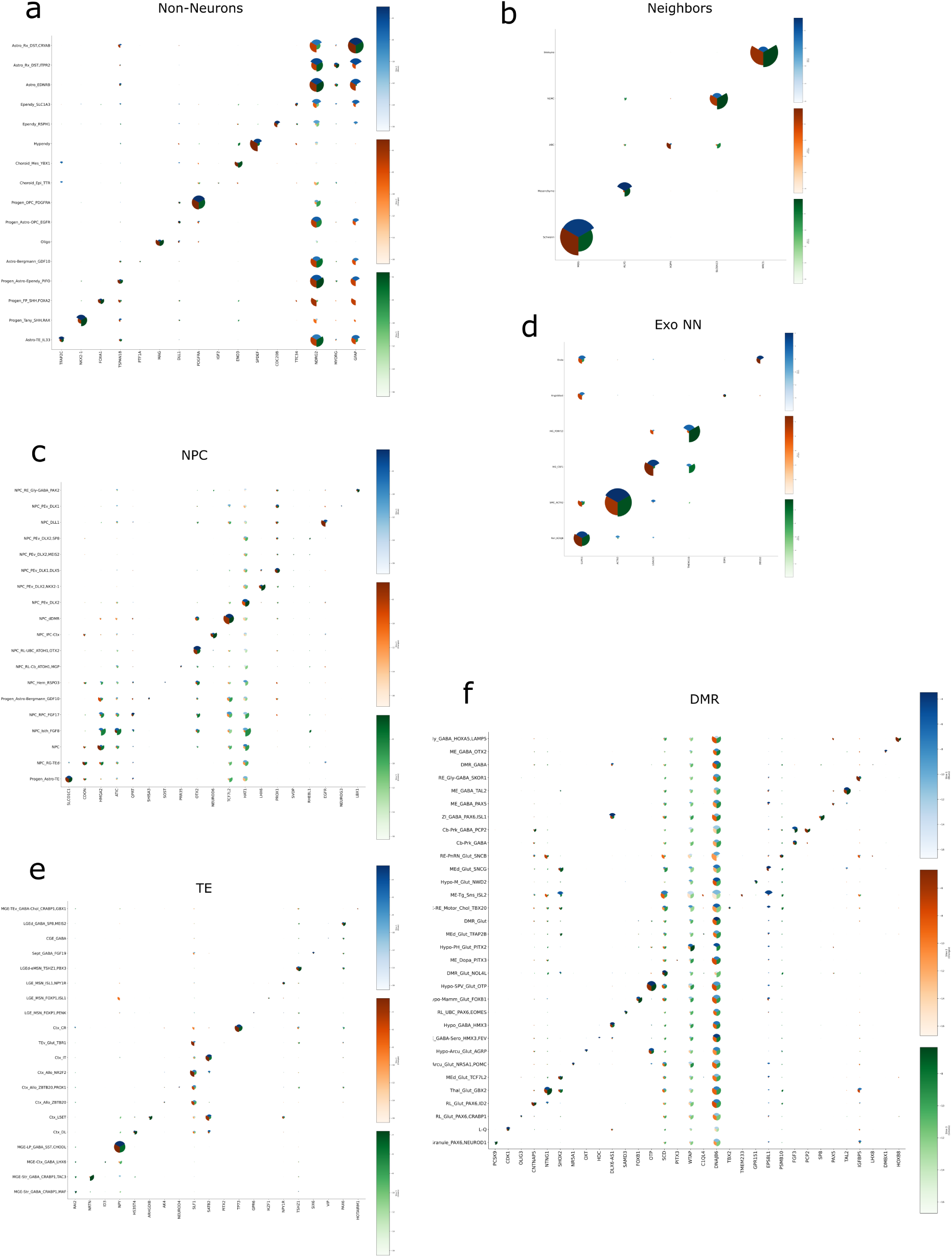
Conserved marker expression across initial classes. Dot plots show mean normalised expression (dot size) and mean inter-species log fold change (color scale; blue = human-up, orange = mouse-up, green = macaque-up) for representative markers of **a** brain-endogenous non-neuronal divisions, **b** brain-adjacent initial classes a.k.a neighbors **c** neuroepithelial / neural progenitor cells (NPC), **d** brain-exogenous non-neurons a.k.a Exo NN, **e** telencephalic (TE) neurons and **f** diencephalic/mesencephalic/rhombencephalic (DMR) neurons.

**Supplementary Figure S4.**
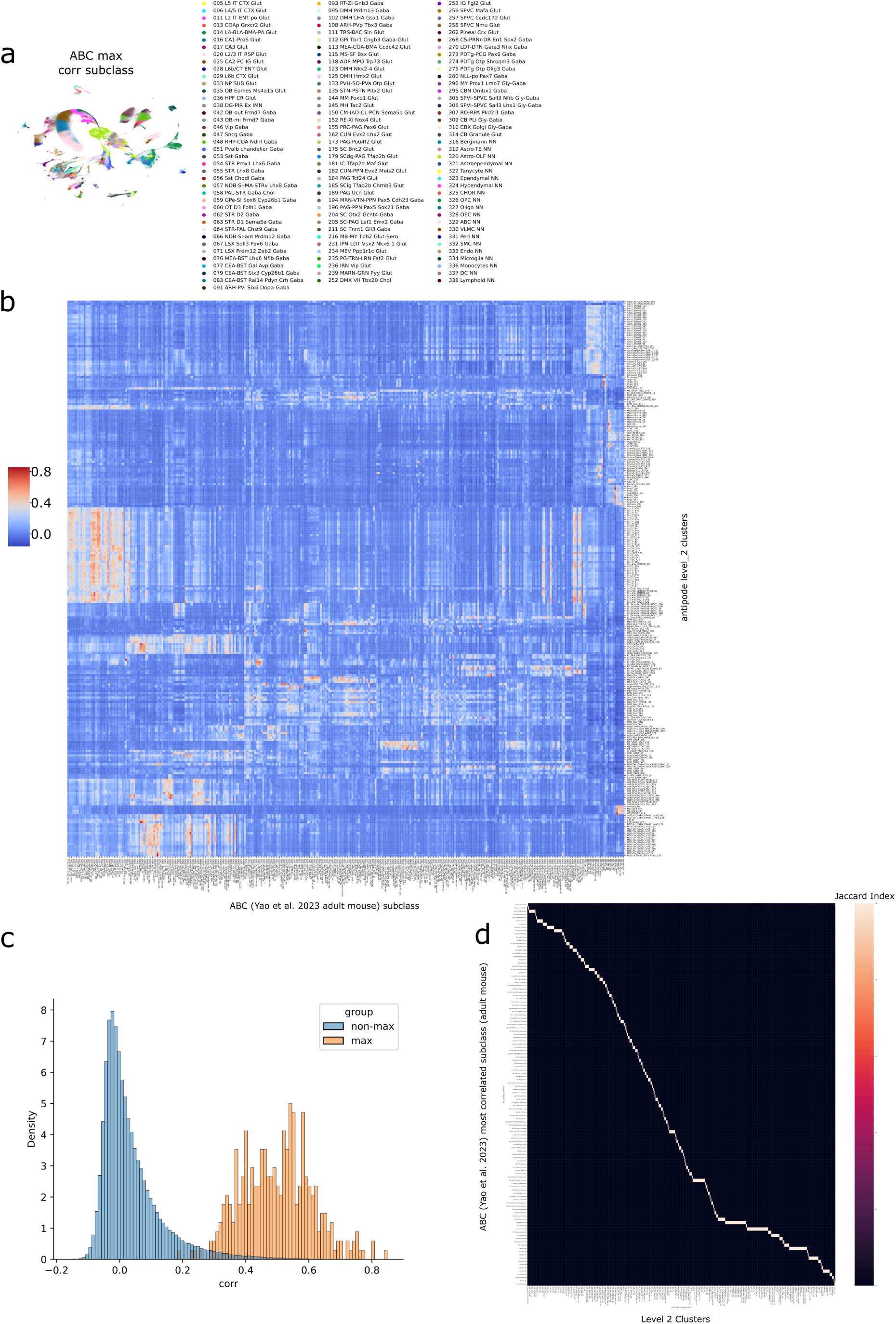
Mapping developing clusters to adult mouse subclasses. **a,** UMAP of developmental clusters colored by their most-correlated adult subclass from Yao et al. 2023. **b,** Heat-map of Pearson correlations (using Yao identity TF set) between 380 developmental clusters (rows) and 337 adult subclasses (columns). **c,** Distribution of correlation coefficients for maximal matches (orange) versus all other comparisons (blue). **d,** Jaccard index of gene-marker overlap for the same developing cluster–adult subclass pairs (because matches are called by cluster all jaccard indices are 0 or 1).

**Supplementary Figure S5.**
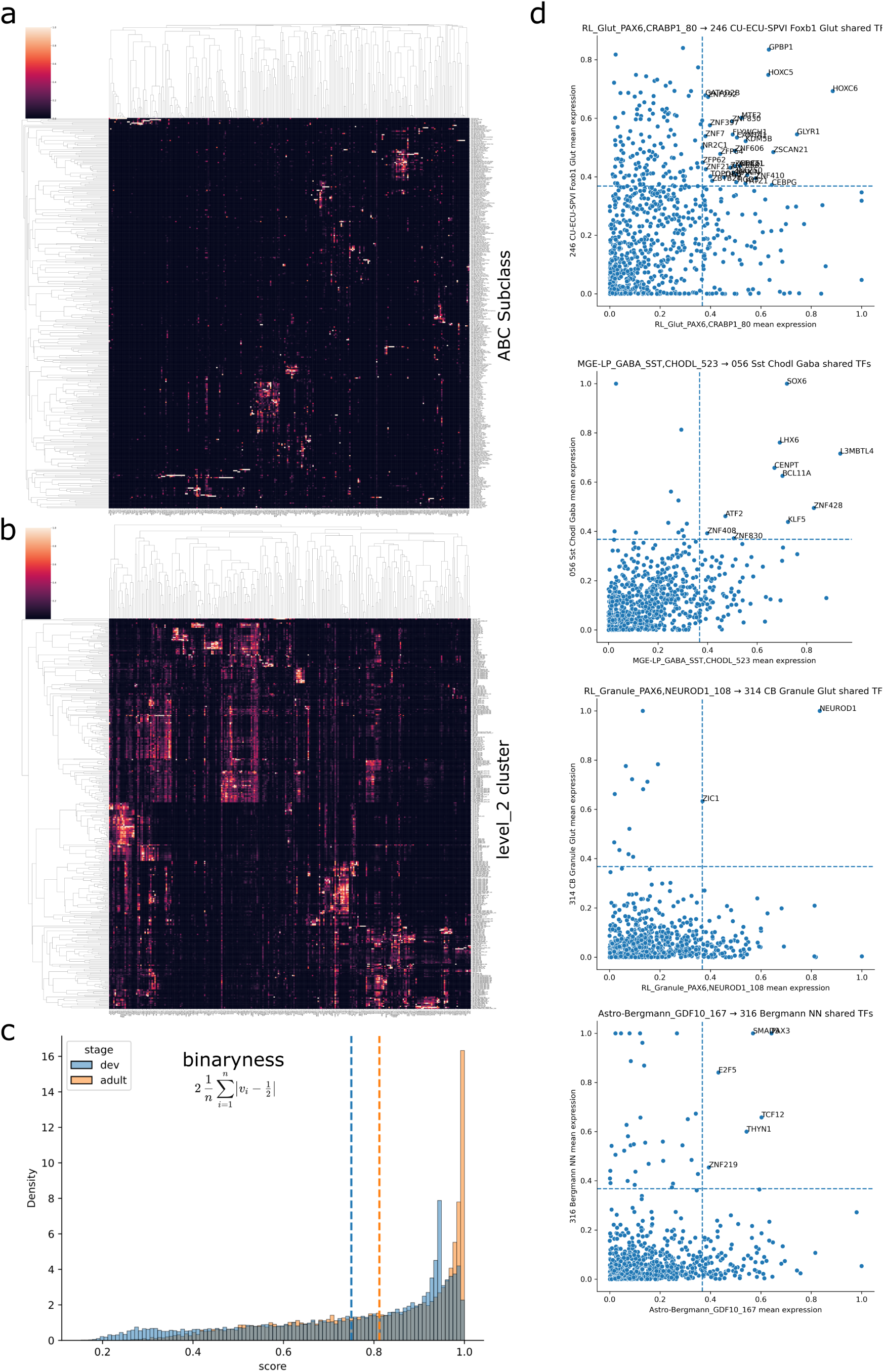
Shared transcription-factor programs between development and adulthood. **a,** Heat-map of binary transcription-factor (TF) expression (presence > 0.1 fraction of cells) across 337 adult cortical subclasses (rows); clustered blocks highlight subclass-specific TF sets. **b,** Equivalent heat-map to **a** for the 380 developmental clusters. **c,** Histogram of TF “bimodality” scores (1 = all-or-none expression) shows higher discreteness in adult (orange) than in development (blue) cell types (dashed lines: medians). **d,** Four example subclass/cluster pairs illustrating TF sharing: each scatter plots mean TF expression in the developmental state (x-axis) versus its matched adult subclass (y-axis); dashed lines mark the 0.33 threshold used to call shared TFs.

**Supplementary Figure S6.**
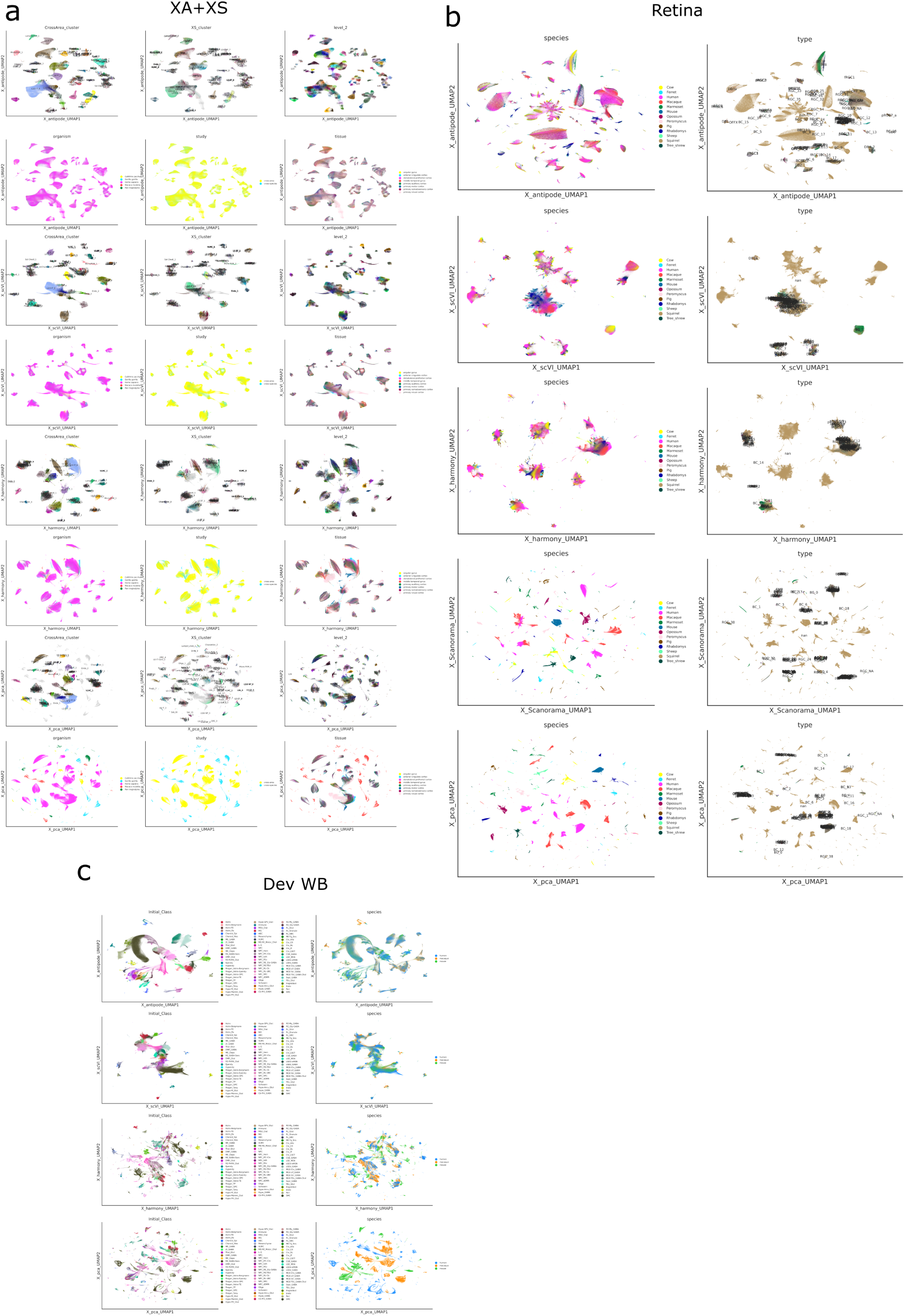
Qualitative comparison of integration methods on three benchmark datasets. UMAPs colored by species (left of each pair) or cell identity (right) for methods used for analysis: ANTIPODE, scVI, Harmony, Scanorama and PCA. **a,** Adult cross-areal (XA) + cross-species (XS) primate cortical dataset integrations. **b,** Adult multi-species mammalian retina. **c,** Developmental whole-brain atlas used in this study.

**Supplementary Figure S7.**
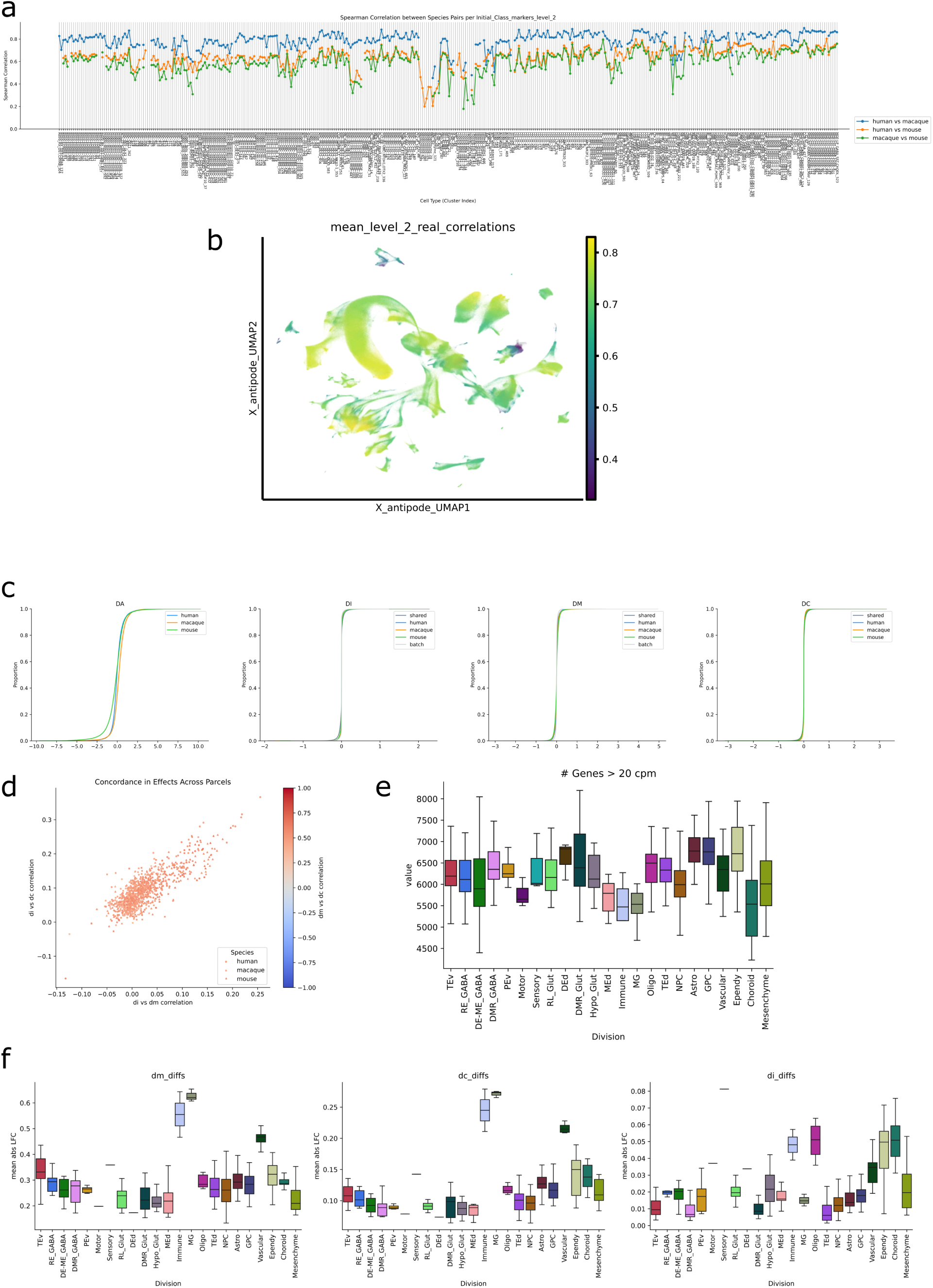
Additional evolutionary gene expression divergence. **a,** Spearman correlation across all species pairs for 380 developmental clusters (named by initial class + cluster). **b,** Mean correlation across species pairs shown in (a) by each cluster, plotted in UMAP space **c,** ECDF plots of raw divergence category parameters. **d,** Scatterplot of the correlation of DM, DC, and DI divergence categories pairwise. Note the consistently high correlation of DC and DM as these are multiplied together in the model (point color). **e,** Boxplot showing the number of genes with mean expression in clusters greater than 20 counts per million (cpm), grouped by division. **f,** Boxplot showing the effect of each divergence category for each cluster, grouped by division. Note that divisions that contain many clusters are biased towards divergence being categorized as DM+DC, while those consisting of fewer clusters seem somewhat biased towards DI.

**Supplementary Figure S8.**
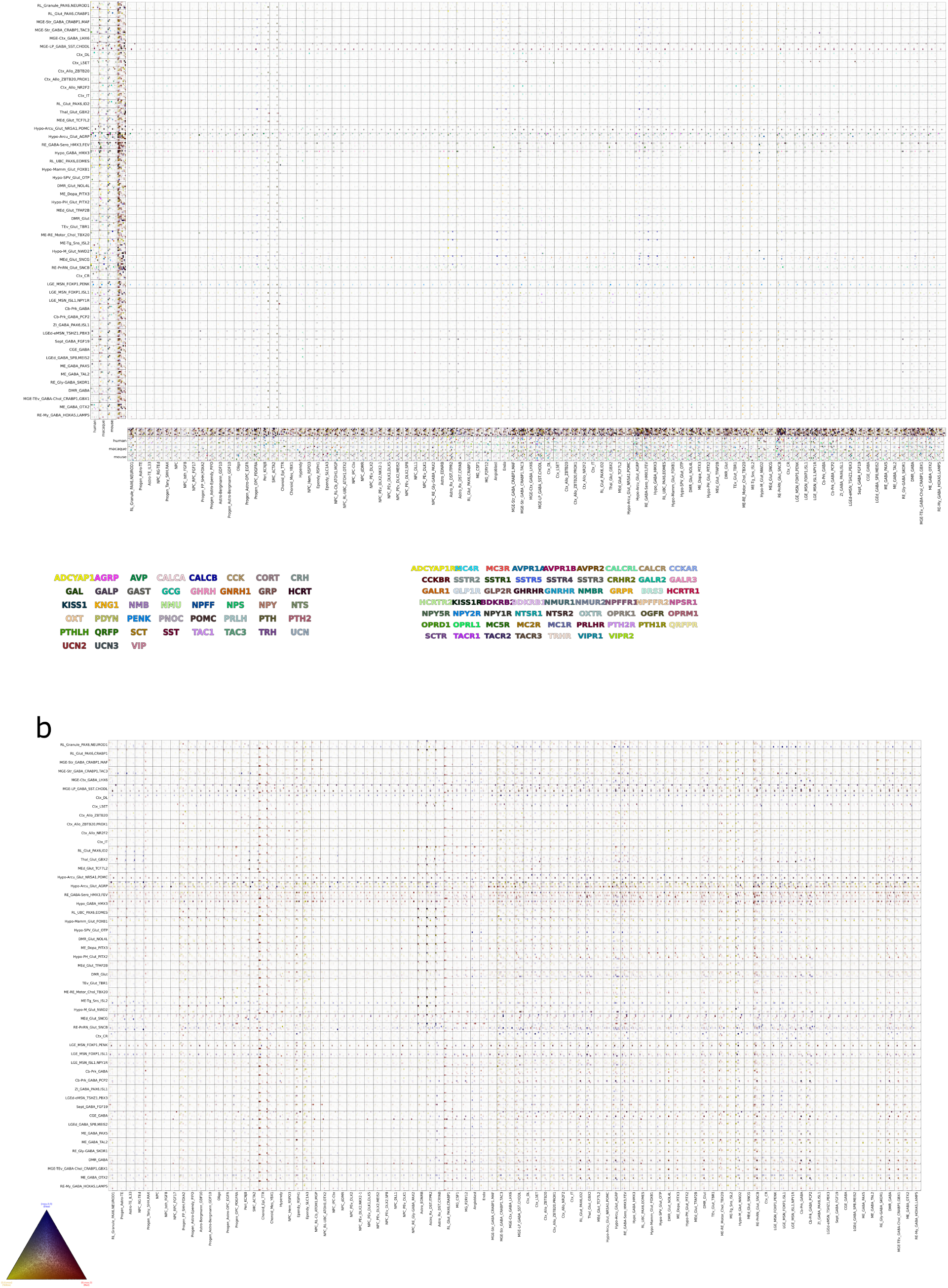
Neuropeptide ligand–receptor expression landscape across progenitor and neuronal states. **a,** Tile-map of neuropeptide-receptor interaction scores (multiplied 0-1 scaled expression). Ligands in neurons (rows) vs receptors in non-neurons (columns). Colored by ligand in main (central heatmap). Along left and bottom shows raw 0-1 expression of ligands/receptors in each species, and tricolor species divergence tile-map. **b,** Tile-map showing species divergence, with 0-1 scaled tri-species means colored according triangular legend.

**Supplementary Figure S9.**
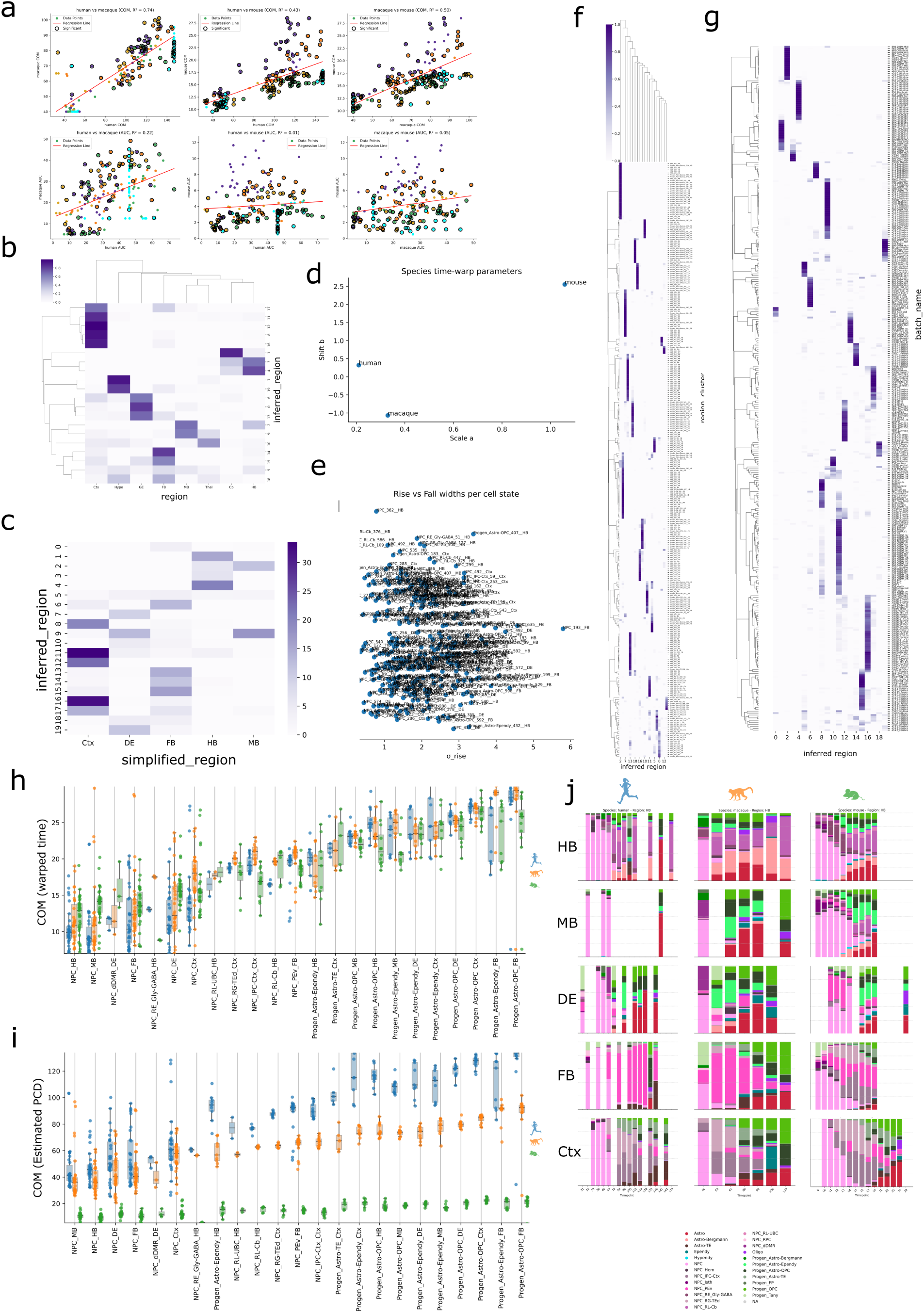
Additional analyses of the progenitor timing model. **a,** Model-free trapezoid sum-based COM calculations based only on raw data for species and regions (colors). **b,** Heat-map of annotated finer regional dissection vs de novo inferred regions. **c,** Heat-map of annotated broad regional dissection vs de novo inferred regions. **d,** Species-level time-warp parameters (scale α, shift Δ) estimated from the model; mouse shows a pronounced left-shift (earlier development). **e,** Global comparison of σₐ versus σᵦ for all progenitor states. **f, g,** Heat-map of ANTIPODE cluster (f) or batch_name (g) vs de novo inferred regions **h, i,** Box plot of fit COM values for each progenitor region + cluster, colored by species in arbitrary warped time (i) or real time (j) and ordered by mean time across species **j,** Stacked barplots showing proportion of total cells included of each progenitor state at each timepoint, for each species, grouped by simplified regions.

## Methods

### Generation of scRNA-seq data

#### Samples

The Primate Center at the University of California, Davis, provided nine specimens of cortical tissue from PCD40, PCD50, PCD65 (n = 3), PCD80 (n = 2), PCD90 and PCD100 macaques. All animal procedures conformed to the requirements of the Animal Welfare Act, and protocols were approved before implementation by the Institutional Animal Care and Use Committee at the University of California, Davis. In total, we analysed single-cell transcriptomes from 654637 cells from developing macaque (this study plus (Zhu et al., 2018)), 473517 cells from developing mouse(Di Bella et al., 2021; Kim et al., 2020; La Manno et al., 2020; Loo et al., 2019; Mayer et al., 2018) and 726613 cells from developing human (this study plus (Bhaduri et al., 2021; Eze et al., 2021; Jessa et al., 2019; Zhong et al., 2023, 2020; Zhou et al., 2022)). De-identified human tissue samples were collected with previous patient consent in strict observance of legal and institutional ethical regulations in accordance with the Declaration of Helsinki. Protocols were approved by the Human Gamete, Embryo, and Stem Cell Research Committee and the Committee on Human Research (institutional review board) at the University of California, San Francisco.

#### Single-cell RNA sequencing tissue processing

For the PCD40 to PCD100 macaques, dissections were performed in PBS under a stereo dissection microscope (Olympus SZ61). Tissue was dissociated and prepared for droplet partitioning as in (Schmitz et al., 2022). Single-cell RNA sequencing was completed using the 10x Genomics Chromium controller and version 2 or 3 3-prime RNA capture kits. Most samples were loaded at approximately 10,000 cells per well; up to 25,000 cells were loaded per lane for a small number of multiplexed samples. Transcriptome library preparation was completed using the associated 10x Genomics RNA library preparation kit. Multiseq barcode library preparation was completed as described in McGinnis et al.46. Following library preparation, libraries were sequenced on Illumina HiSeq and NovaSeq platforms.

#### Alignments and gene models

Fastq files were generated from Illumina BCL files using bcl2fastq2. Genes were quantified using Kallisto release 0.46 using each species respective reference: the human hg38 genome assembly and gencode version 33 transcript annotation, the RheMac10 genome assembly, annotated using the comparative annotation toolkit based on the human annotation48, and the transcript annotations of Mus musculus ENSEMBL release 100.(Bray et al., 2016) A custom Kallisto reference for each species was created for the quantification of exons and introns together, in which introns were defined as the complement of exonic and intergenic space. The Kallisto index used k-mers of length 31. Public data were downloaded as raw fastq files or as BAM files that were converted back to fastq files. All data were processed from raw reads using the same Kallisto pipeline to minimize annotation and alignment artefacts. 16738 1:1:1 ortholog genes were identified using the MGI human-mouse ortholog table, and this geneset was used for analysis.

#### Quality control

Kallisto–Bus output matrix files (including both introns and exons) were input to Cellbender (release 0.2.0; https://github.com/broadinstitute/CellBender), which was used to remove probable ambient RNA only.(Fleming et al., 2023) Only droplets with a greater than 0.99 probability of being cells (not empty droplets), as calculated by the Cellbender model, were included in further analysis. Only spliced read UMIs were used thenceforth. Droplets with fewer than 800 genes detected, or greater than 40% ribosomal or 15% mitochondrial reads, were filtered from the dataset. Finally doublets were detected using solo and cells with a doublet probability greater than 0.4 were excluded. Non-brain cell types (neural crest, placode, etc.), undetected doublet clusters and low quality cells were removed from the dataset by author judgement.

### Processing of data

Raw fastq files were obtained from the NCBI Sequence Read Archive or directly from authors. Transcriptomic profiles were quantified using kallisto 0.46.0, with references built using gencode v33, comparative annotation toolkit rheMac10 (based on human gencode v33) and mm10 for human, macaque and mouse respectively. Exonic and intronic counts were then passed to cellbender using only the remove ambient model with parameters.

The ANTIPODE model is described in supplement 1. The data analysis herein uses the model trained with 600 starting clusters, 100 latent dimensions, a Laplace dispersion prior on posterior parameter estimates of 500 an encoder with layers of size [#genes, 6000,3000,100], and a classifier with layers of size [100,3000,3000,600].

### Calculation of gene expression estimates

Direct calculation of gene expression means in clusters was formulated as the sum of counts for a gene across the cells assigned to a taxon divided by the total UMI counts, where a Laplace pseudocount is defined as [inline] is added. The division by 2 stems from the naive estimate of the number of additional counts required to observe a count for a gene for which no counts have been observed following the memorylessness property of the geometric distribution. ANTIPODE parameter-based calculation of gene expression of means [inline]. Marginal gene expression differences attributed to DA, DM, DI and DC are calculated as [inline], where [inline] is the mean calculation without the respective parameter. *L* represents cluster gaussian samples in latent space, *DM_c,s_* is the differential-by-module with respect to cluster and species, *W* is the matrix of latent space components to genes weights, *DC_s_* represents the differential-by-coexpression term with respect to species, *DI_c,s_* represents the differential-by-identity intercept with respect to species and cluster and *DA_s_* represents the differential-by-all intercept with respect to species only. In the model fitting, terms for batch effects by module, batch effects by identity, and secondary covariate by module effects are added as well (see code and Supplement 1).

### Lateness

Regional progenitor taxa were defined as antipode clusters split by the simplified region (pallium, ventral forebrain (FB), diencephalon (DE), mesencephalon (ME) and rhombencephalon) for the neuroepithelial NPC initial class or simply antipode clusters for all other initial classes, as the NPC states tended to be similar enough to be undifferentiated by the model across regions. Regional progenitor taxon proportions (RPTP) were calculated as the number of cells from a droplet based sequencing batch divided by the total number of progenitors and endogenous glia (progenitors and glia excluding microglia). Raw lateness values for comparison were calculated as the center of mass of RPTP of the curve RPTP = f(Timepoint), calculated by the trapezoid sum. The lateness Bayesian models for lateness COM estimates in the whole brain and cortical areas are pyro models, the code for which can be found in the project github repository under analysis 5-2.

### Neuropeptide ligand-receptor interactions

We calculated the interaction score following the method used by NATMI (Hou et al., 2020), where the score for each ligand is equal to the mean of the ligand expression multiplied by each potential receptor’s expression. Expression values used were the true counts/count value in each initial class normalized by the maximum expression of that gene in any initial class. The concordance of receptor and ligand expression divergence was calculated by the, compared to the null distribution of this value calculated from permuted receptors and ligand values across initial classes.

### Other bioinformatic analysis

Disease enrichments were performed using the gseapy package’s preranked gene list test with FDR q values calculated with at least 1000 permutations. Gene ontology enrichments vs lateness score means were computed using permutation tests. The whole dataset UMAP layout was chosen from 100 random seeds of the cuML accelerated UMAP algorithm, with the displayed layout chosen based on the amount of space surrounding the DMR neuron classes for labeling. Mapping of adult subclasses to developing initial classes was based on the maximum Pearson correlation of counts per count values scaled from 0-1 per gene, using the list of cell type identity TFs provided in Yao et al.(Yao et al., 2023).

### Large language model statement

LLMs were used to proofread and point out areas for textual clarity improvement, however none of the manuscript text was generated by an LLM. The ANTIPODE model was written without aid from an LLM, however later iterations and code organization and documentation was aided by ChatGPT. Some code to perform analysis for figures was written or reorganized by LLM.

## Supplementary Tables

Supplement 1. Specification and further discussion of the ANTIPODE generative model.

Supplementary Table 1. Metadata for each sample in the meta-atlas, including the species, closest region annotation, developmental time point, individual and sequencing quality-control metrics

Supplementary Table 2 Dictionary of initial classes. Qualitative definitions of classes explored in the atlas with extended explanations for inferences about initial–terminal class relationships.

Supplementary Table 3 Gene scores for the various lateness metrics

## Acknowledgements

We thank Alice Tarantal for providing samples, Min Cheol Kim for discussions related to the ANTIPODE model, and Anna Wright for editing assistance.

We acknowledge the following funding sources: Ruth L. Kirschstein National Research Service Predoctoral Fellowship Award F31 F31NS124333 (M.T.S.), DP2MH122400-01 (A.A.P.), U01MH114825 and UM1MH130981 (A.A.P., T.J.N.), R01AI136972 (C.J.Y.), R01MH134981 (A.A.P.) the Chan Zuckerberg Biohub (C.J.Y., T.J.N., A.A.P.), National Institutes of Health DP2MH122400-01, Schmidt Futures Foundation, Shurl and Kay Curci Foundation Innovative, W.M. Keck Foundation, and William K. Bowes Jr. Foundation. A.A.P. and T.J.N. are New York Stem Cell Foundation Robertson Investigators and members of the UCSF Kavli Institute for Fundamental Neuroscience.

## Data availability

The sequencing data have been deposited in the Gene Expression Omnibus under accession number GSE306257; the data are browsable at dev-whole-brain-hqm.cells.ucsc.edu. Scripts and annotation files for the study have been deposited on github at https://github.com/mtvector/antipode_manuscript and the ANTIPODE package can be found at https://github.com/mtvector/scANTIPODE.

