## Supplementary Document 1 for "ANTIPODE Provides a Global View of Cell Type Homology and Transcriptomic Divergence in the Developing Mammalian Brain"

### ANTIPODE: an evolution-inspired generative model of cell types

Matthew Schmitz

#### Single Cell Ancestral Node Taxonomy Inference by Partitioning Of Differential Expression

ANTIPODE follows the paradigm of SCVI [Lop+18] and its child scANVI[Xu+21], being a probabilistic variational autoencoder and a generative model of single cell sequencing data. The main philosophy behind the development of ANTIPODE is that integration of single cell data can be thought of as simultaneously clustering data while performing differential expression/coexpression, and that integration along biological axes of variation can be achieved by accounting for known forms of gene expression modulation. In order to make learned parameters more biologically interpretable, however, ANTIPODE jettisoned elements that make SCVI models more robustly trainable, specifically multiple layer nonlinear decoding, unstructured latent space and decoder batch normalization.

ANTIPODE finds ways around these challenges with stepwise initialization, while also imposing a stricter generative structure within the latent space. The latent space is decoded in a single layer, imposing Laplace distribution priors on parameters to learn a parsimonious model fit of the core parameters composing classical differential expression: differential by module $DM$, differential by coexpression $DC$ and differential by identity $DI$. This is similar in motivation to the LD-VAE to attempt to make SCVI more interpretable, however the extra terms for $DC$ and $DI$ increase flexibility[Sve+20], and the addition of a structured generative model further increases interpretability. ANTIPODE's model is built purely within pyro[Bin+19] which provides the mathematical Bayesian framework and inference engine.

Similar to SCVI followed by scANVI model learning, ANTIPODE is trained in multiple phases. In **phase 1** (the *fuzzy phase*), the $RelaxedCategorical$ distributions are replaced by $RelaxedBernoulli$s, learning a preliminary version of the model with fuzzy clustering (cells can belong to many clusters at once). The fuzzy phase was an accidental discovery and greatly improves the trainability of the model, as the initialization of ANTIPODE in a state which doesn't explode is not trivial. In **phase 2** (which can be a standalone supervised model if clustering and/or a latent space is provided) supervised clusters are initialized and a constrained version of the model below is fit (Samples from (6) and/or (8) below are treated as observed). Finally in **phase 3**, the model trains without constraint using the generative structure shown in figure 1.

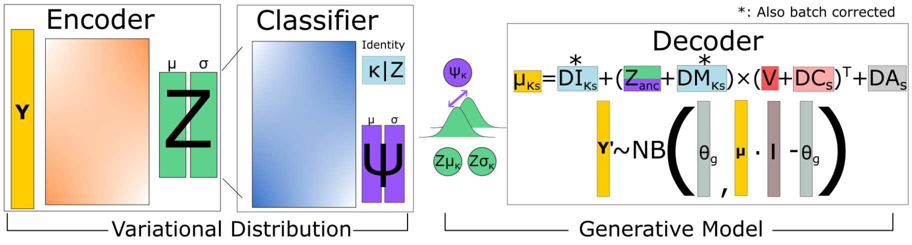


Figure 1: Schematic of the ANTIPODE generative model and variational guide

#### Variables Definition

$Y$ - Gene expression counts.

$Y'$ - Reconstructed gene expression counts.

$G$ - Number of variables (genes).

$Z$ - The latent space.

$V$ - The module $\times$ genes weight matrix (the $Z$ decoder weights).

$Z_{loc}$ - The cluster $\times$ module gaussian centers in latent space.

$Z_{\sigma}$ - The cluster $\times$ module gaussian scales in latent space.

$I_{\kappa}$ - Identity $\times$ gene effects for cluster $\kappa$.

$q$ - A [0,1] quality score affecting the dispersion of the reconstruction $NegativeBinomial$.
note: $Q$ and $q$ can be toggled off in the model if quality isn't a concern or q is being learned inappropriately.

$Q$ - 1D vector of (1-$q$) effects on each gene.

$\beta$ - Batch (b) effects on modules or intercepts. Note for $\beta I$: b $\times$ K $\times$ g is too many parameters, so b is replaced by a batch embedding.

$D$ - Species (s) effects for $DM$ modules, $DC$ coexpression, and $DI$ identity, or $DA$ constitutive species/annotation effects. Note that $DA$ is not sampled from the Laplace distribution and is a free parameter, allowing it to serve as an intercept to account for genes which may be altogether missing from species.

$\phi$ - The module dynamics effected upon $Z$. note: $\phi$ and $\psi$ can be toggled on/off in the model if cell types are believed to be discrete.

$\psi$ - A [-1,1] within-cluster "pseudotime" value multiplied by $\phi$.

$\alpha$ - Probability of cell in each cluster, simplex in phases 2,3.

$\kappa$ - The discrete cluster a cell belongs to.

$Z_\mu$ - [Variational estimation of] the cell's likely mean $Z$ location.

$\hat{I}$ - Adjusted identity effects considering $DI_{s\kappa}$, batch effects $\beta_{b\kappa}$, species effects $D_s$, and modified $Q$, $\hat{Q}$.

$\hat{\mu}_{\kappa sb}$ - The reconstructed gene expression for a given cluster + species + batch.

$\ell$ - The scale factor (sum of counts per cell). Note: SCVI treats this as learnable, I do not.

$\theta$ - Per gene inverse dispersion for final negative binomial (the total failures parameter commonly called $r$).

$\hat{\mu}_\kappa$ - Estimated gene expression in cluster $\kappa$ without batch or species effects.

### ANTIPODE Generative Model

$$V, Z_{loc}, I_\kappa, Q, \beta, D, \phi, \psi \sim Laplace\,(0, priorscale) \tag{1}$$

$$Z_\sigma \sim HalfCauchy\,(1) \tag{2}$$

$$q \sim expit\,(Logistic\,(0,1)) \tag{3}$$

$$\hat{Q} = (1-q)\cdot Q \tag{4}$$

$$\alpha \sim Dirichlet\,(1,1,...) \tag{5}$$

$$\kappa \sim RelaxedCategorical\,(\alpha_{0...K}) \tag{6}$$

$$Z_\mu = Z_{loc_\kappa} + ((2\cdot expit(\psi)-1)\cdot\phi_\kappa) \tag{7}$$

$$Z \sim Normal\,(Z_\mu, Z_\sigma) \tag{8}$$

$$\hat{I} = I_\kappa + DI_{s\kappa} + \beta_{b\kappa} + D_s + \hat{Q} \tag{9}$$

$$\hat{\mu}_{\kappa sb} = softmax\left((Z + DM_{s\kappa} + \beta_{b\kappa})(V + DC_s)^T + \hat{I}\right) \tag{10}$$

$$\ell = \sum_{j=0}^{G} Y_{ij} \tag{11}$$

$$\mu_{\kappa sb} = log(\ell \cdot \hat{\mu}_{\kappa sb}) \tag{12}$$

$$\hat{\theta} = q \cdot \theta \tag{13}$$

$$Y' \sim NegativeBinomial\left(\hat{\theta}, expit\left(\mu_{\kappa sb} - log(\hat{\theta})\right)\right) \tag{14}$$

As such, the estimated gene expression in a cluster $\kappa$ is given by:

$$\hat{\mu}_{\kappa} = Z_{\mu\kappa} V^T + I_{\kappa} \tag{15}$$

### ANTIPODE Variational Guide

$$V, Z_{loc}, Z_{\sigma}, I_{\kappa}, \beta, D, \phi, \psi, Q \sim Delta\left(V, Z_{loc}, Z_{\sigma}, I_{\kappa}, \beta, D, \phi, \psi, Q\right) \tag{1}$$

$$Z_{\mu}, Z_{\sigma}, q_{\mu}, q_{\sigma} = f(Y, s) \tag{2}$$

$$Z \sim Normal\left(Z_{\mu}, Z_{\sigma}\right) \tag{3}$$

$$q \sim Normal\left(q_{\mu}, q_{\sigma}\right) \tag{4}$$

$$\alpha, \psi_{\mu}, \psi_{\sigma} = g(Z) \tag{5}$$

$$\psi \sim Normal\left(\psi_{\mu}, \psi_{\sigma}\right) \tag{6}$$

$$\kappa \sim RelaxedCategorical\left(\alpha\right) \tag{7}$$

$f(x)$, $g(x)$ represents the encoder and classifier deep neural networks, respectively. Equation 1 is meant to show that all of these parameters are passed directly to the model by sampling them from a *Delta* distribution (how MAP inference is achieved in pyro).

I note that the model contains a few extra features, including the ability to learn a layered tree for a hierarchical $\kappa$, in which the leaf clusters are propagated up the tree by cross-multiplying edge weight matrices pseudo-discretized by *RelaxedCategorical* sampling. In addition, instead of providing species as a single onehot value, the model allows for any structure of discovery covariate design matrices, allowing more complex (soon extending to phylogenetic) regression.

With priors and point estimates of the parameters, the model gives a maximum a posteriori parameter estimate for most parameters. Calculation of the ELBO is performed automatically by the pyro probabilistic programming language[Bin+19], and parameter optimization is performed by the Adam optimizer using the ELBO as a loss function. Note: Pyro's stochastic variational inference (SVI) class was modified (SafeSVI) to track a moving window of the ELBO and if this value is infinite or more than N standard deviations from the moving average, the optimizer is prevented from taking a step.

### Additional Information

#### Layered Tree Clustering

Rather than learning a flat clustering, the model is capable of learning (or taking supervised input) for a layered tree with $N$ layers of the format layer sizes = $[L_0, L_1 ... L_N]$ where $L_0 = 1$ and $L_{n+1} > L_k$, representing the number of clusters at each level (see Figure 2). In phases 2 and 3, $\kappa$ is sampled from a *RelaxedCategorical* distribution yielding a matrix one hot row vectors of size for each cell. In order to get the membership layered tree, we will multiply by the adjacency matrix of a bipartite graph between each sequential layers of the tree. For these adjacency matrices, we first sample tree edge weights from a *RelaxedCategorical* (edge weight matrices will be size $[L_n, L_{n-1}]$) and so each $\kappa$ in $L_n$ belongs to only one $\kappa$ in $L_{n-1}$. Therefore, after propagating the $\kappa$ up the tree, each cell's $\kappa_L$, the membership across all $N$ layers, is $\sum_{l=0}^{N} \kappa_l = N$.

#### Parameter Sampling Scaling

As many of these parameters' number of sites sampled scales with minibatch size $\times$ latent dimension $\times$ number of genes, it was necessary to sample them efficiently as to not overflow hardware limitations. Resampling the same parameters the number of times in each minibatch increases computation beyond VRAM capacities and simply sampling once per parameter: numper of species $\times$ latent dimension $\times$ number of gene penalizes rarer types/dimensions/species. *pyro.poutine.scale* provides a utility for scaling the ELBO contribution of sites within plates. For this purpose I developed a function, flexible einsum scale tensor ($fest(x, y, z)$) which multiplies out the marginal of the design matrices, or $\kappa$ onehot vector/$Z$ component absolute value, to get the normalized joint of any number of vectors, which scales each parameter as if it were sampled once per gene, in proportion to the number of times a site would have been sampled, for instance if $DI$ were sampled the number of times each species and $\kappa$ occurred

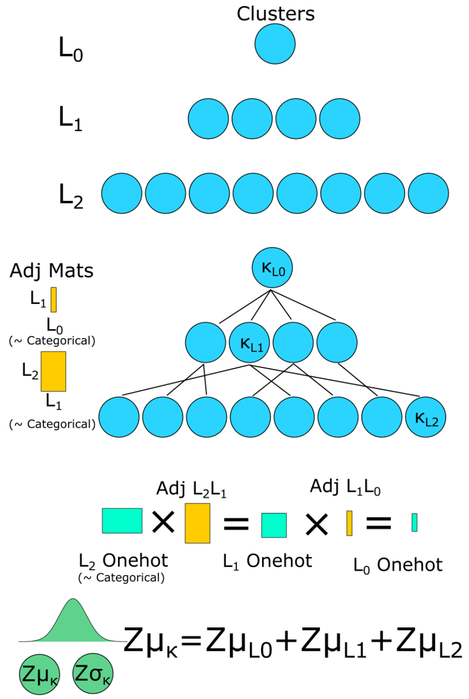


Figure 2: The process for generating optional hierarchical clustering in ANTIPODE. Note that the adjacencies and the bottom layer are sampled from OneHotCategorical distributions, such that one cell belongs to one leaf cluster, and each child cluster belongs to one and only one parent.

in the minibatch. Thus the sum scale for the elbo at each site is batch size $\times$ $G$ (or the number of latent dimensions). This scaling is critical to the trainability of the model.

### Continuing Challenges

#### Phylogenetic Regression

This model is designed (see the A-N- in ANTIPODE) to be capable of learning the ancestral latent space by inclusion of phylogenetic contrasts in the discov (originally discrete covariate, but now discovery covariate) and prior scales. Research into the correct way to implement this is ongoing. Please reach out if you would like to offer assistance on this point.

#### Parameter Variances/Confidence

ANTIPODE is currently unable to estimate the variance of its learned parameters of interest and thus is mostly a MAP model rather than a Bayesian one. This means that currently only point estmates of parameters, which makes determination of significance of differences a matter of ranks rather than per-parameter confidence intervals or p-values. It is unclear if implementation of Bayesian parameter sampling modules will be possible as the model is too large to apply Markov Chain Monte Carlo methods, variational inference tends to struggle with properly estimating, and the proper allocation of variance between the reconstructed observed $NegativeBinomial$, cluster $Normal$ distribution, and parameter confidence is currently an unsolved problem.
